# Clinal variation in drought response is consistent across life stages but not between native and non-native ranges

**DOI:** 10.1101/2023.09.28.559709

**Authors:** Dávid U. Nagy, Arpad E. Thoma, Mohammad Al-Gharaibeh, Ragan M. Callaway, S. Luke Flory, Lauren J. Frazee, Matthias Hartmann, Isabell Hensen, Kateřina Jandová, Damase P. Khasa, Ylva Lekberg, Robert W. Pal, Ioulietta Samartza, Manzoor A. Shah, Min Sheng, Mandy Slate, Claudia Stein, Tomonori Tsunoda, Christoph Rosche

## Abstract

Clinal variation, i.e., intraspecific variation that corresponds to environmental gradients, is common in widely distributed species. Studies on clinal variation across multiple ranges and life stages are lacking, but can enhance our understanding of specieś adaptive potential to abiotic environments and may aid in predicting future species distributions.

This study examined clinal variation in drought responses of 59 *Conyza canadensis* populations across large aridity gradients from the native and non-native ranges in three greenhouse studies. Experimental drought was applied to recruitment, juveniles, and adult stages.

Drought reduced growth at all three life stages. However, contrasting patterns of clinal variation emerged between the two ranges. Native populations from xeric habitats were less inhibited by drought than mesic populations, but such clinal variation was not apparent for non-native populations. These range-specific patterns of clinal variation were consistent across the life stages.

The experiments suggest that invaders may succeed without complete local adaptation to their new abiotic environments, and that long-established invaders may still be evolving to the abiotic environment. These findings may explain lag times in some invasions and raise concern about future expansions.

## Introduction

Water availability is a major determinant of plant growth, reproduction, abundance, and distribution. Plants deal with low water availability through plastic and/or adaptive strategies (Volaire 2018), for example, (1) plants can escape dry periods by shortening their life cycle and altering their phenology or (2) they can avoid drought by adjusting functional traits related to transpiration (e.g., specific leaf area, root shoot ratio, leaf dry matter content). Surprisingly, the extent to which functional traits can predict plant performance in response to drought remains unclear (Westerband et al 2019; González de Andrés et al. 2021). This is a major issue in global change ecology because the frequency and severity of drought events are rapidly shifting through climate change (Cook et al. 2018). In particular, the understanding of adaptation to drought is crucial for predicting future species distributions, such as range expansions by invasive species or the persistence of plant populations experiencing climate change (Moritz 1994; Palkovacs et al. 2012; Pratt & Mooney 2013; Broennimann et al. 2014; Exposito-Alonso et al., 2019).

Investigating clinal variation is an effective approach for studying the adaptive potential of plants, i.e., studying intraspecific variability in the response to an experimental stress using populations that occur along environmental gradients (Montesinos-Navarro et al. 2010; Alexander et al. 2012; Pratt & Mooney 2013). There are several possible patterns when comparing the performance of populations originating from a gradient in aridity (mesic to xeric) when grown under dry vs. wet experimental treatments (Fig. 1). Plants may perform the same across the entire gradient (i.e., no evidence for adaptation) or the same in wet vs. dry conditions (i.e., no treatment effect / no plasticity, Fig. 1a). Alternatively, plants may grow less in dry treatments than wet treatments but do so in the same way across the aridity gradient (i.e., no clinal variation / evidence for adaptation to increasing drought, Fig. 1b). Clinal variation would be evident if plants from mesic habitats demonstrate a large positive change in performance between wet vs. dry conditions (ΔP) whereas plants from xeric habitats showed either a small positive ΔP or a negative ΔP (i.e., plants from xeric habitats perform better in dry treatments than in wet treatments, Fig. 1c or Fig. S1 in the Supplements for more details). This would create a significant two-way interaction between response to experimental drought and mesic vs. xeric population sources. The opposite pattern may occur when ΔP increases with increasingly xeric habitats (Fig. 1d). However, this pattern would be unexpected as populations from xeric environments would appear to be less adapted to drought than populations from mesic environments.

**Figure 1.**
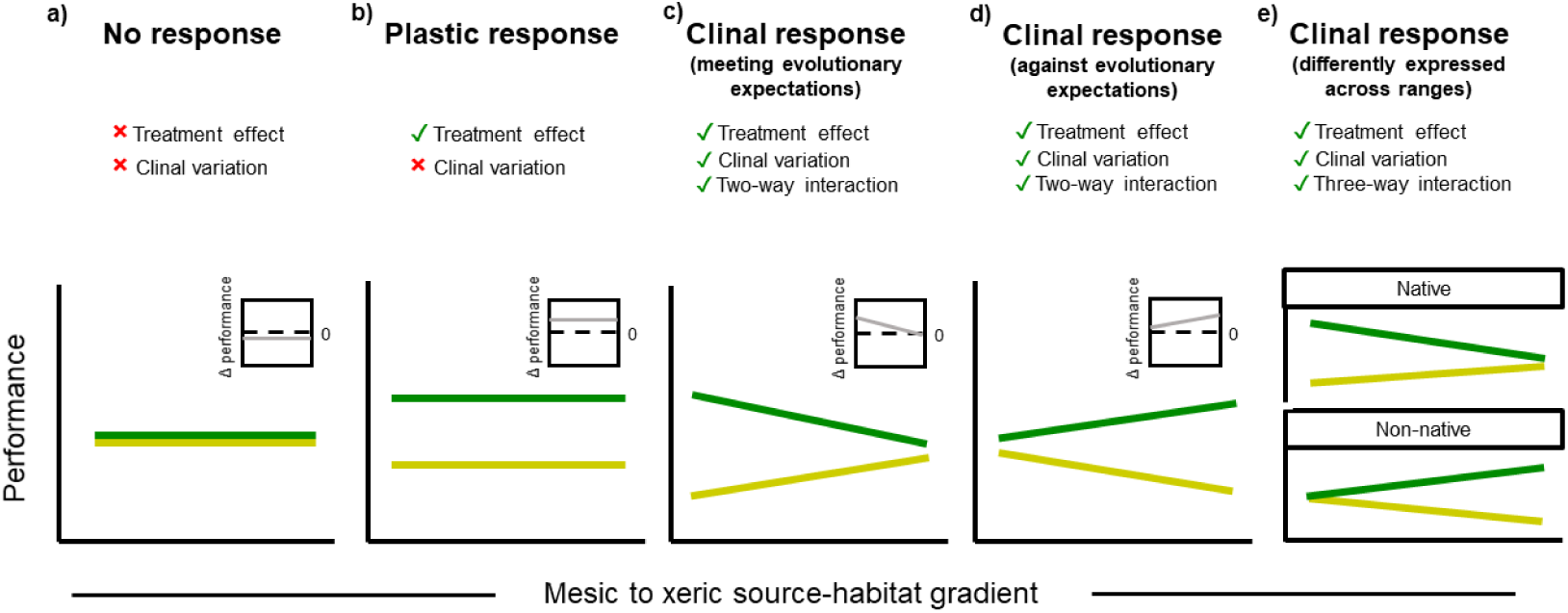
Predictions for the performance of plant populations under wet (green lines) and dry experimental conditions (yellow lines) along an aridity gradient in how often and severe drought occurs in their population history (i.e., aridity gradient from mesic to xeric source-habitat). For the sake of clarity, the populations have not been included (e.g., as dots) but only their linear response slope for the two drought treatments along their aridity gradient. In a) performance does not respond to the experimental drought and shows no signs of clinal variation along the drought gradient. In b) experimental drought affects plant performance (Δ performance), but the drought gradient does not. In c) and d) applied experimental drought affects plant performance and the drought gradient affects Δ performance (i.e., clinal variation can be explained by the drought gradient). Out of the two adaptive responses c) meets evolutionary expectations where increasing drought gradient in the population history results in decreasing Δ performance, while d) presents the opposite response, i.e., against evolutionary expectations. In e) the differently expressed clinal variation between native and non-native range is presented. A more detailed figure including populations and different estimated Δ performances along the gradient can be found in Supplementary data (S1).

Studying broad spatial and bioclimatic gradients in clinal variation in response to water supply can estimate the adaptive potential of species, but only a few studies have sampled broad geographic gradients (Cook et al. 2018). Among these, Exposito-Alonso et al. (2019) examined the effects of experimental drought on 517 *Arabidopsis thaliana* populations. They showed that populations from the edges of the environmental limits of *A. thaliana* experienced the strongest climate-driven selection. However, Exposito-Alonso et al. (2019) included only native Eurasian and North African populations in their experiment, and these populations represent a very long history of *in situ* evolution (i.e., long population history in aridity).

In contrast, comparing mesic to xeric gradients for native and non-native populations allows exploration of natural selection at two very different evolutionary time scales, and this might provide insight into the speed at which selection might occur. Non-native populations often experience rapid range expansions across broad environmental conditions (Broennimann et al., 2014) which can lead to more random distributions of genotypes in non-native ranges than in native ranges (Keller & Taylor 2008; Rosche et al. 2016; 2018; Nagy et al. 2018). Non-native populations may counteract this through rapid evolution (Callaway & Maron 2006; Vandepitte et al. 2014; van Kleunen et al. 2018). However, whether rapid evolution leads to comparable clinal variation in drought responses between native vs. non-native ranges is poorly understood. Mráz et al. (2014) and Villellas et al. (2021) found that clinal variation to dry and wet experimental treatments was more pronounced in native than in non-native populations of *Centaurea stoebe* and *Plantago lanceolata*, respectively. Instead, non-native populations of *P. lanceolata* responded with more plasticity to experimental drought than native populations (Villellas et al. 2021), which might be due to the lack of time to adapt to the new environments. In the context of Figure 1, such differences in clinal variation would be illustrated by a significant three-way interaction (range × experimental drought × aridity clines, Fig. 1e).

The few mentioned studies that sampled broad bioclimatic gradients focused exclusively on adult plants to quantify clinal variation in drought responses (e.g., Mráz et al. 2014; Exposito-Alonso et al. 2019; Villellas et al. 2021). On smaller bioclimatic scales, there are studies that investigated clinal variation at early life stages (e.g., Al-Gharaibeh et al. 2017; Elnaggar et al. 2019). However, we are not aware on studies that compared differences in drought responses across several plant life stages. This comparison may be important because plants show many distinct ontogenically-based traits and drought can have far stronger effects on seedling performance than on adults (Niinemets 2010; Coe et al. 2021). Thus, selection pressure should vary at different life stages (Mitchell & Bakker 2014; Zirbel & Brudvig 2020; Havrilla et al. 2021). Shifts in the timing of drought events – as predicted through climate change – could be particularly detrimental at early life stage that are less experienced with drought (e.g., Parmesan & Hanley 2015; D’Orangeville et al. 2018).

To address the apparent research gaps, this work studied clinal variation in drought response for 30 native and 29 non-native populations of *Conyza canadensis* (Canadian fleabane) collected across broad aridity gradients in both the native and the non-native ranges. In a greenhouse, three complementary experiments were conducted in which (i) recruitment (i.e., germination and early seedling development), (ii) juvenile, and (iii) adult life stages to wet and dry treatments were subjected. Predicting that:

(1) Plant performance under experimental drought correlates with functional traits related to transpiration (specific leaf area, root shoot ratio, leaf dry matter content) because these traits mediate drought tolerance.
(2) Due to local adaptation, populations from xeric habitats exhibit greater tolerance to experimental drought than populations from mesic habitats, resulting in clinal variation (Fig. 1c).
(3) Because of a longer timeframe for evolution, clinal variation in response to wet-dry treatments is more pronounced in native populations than in non-native populations (i.e., three-way interactions between aridity in population history, range, and experimental drought, Fig. 1e).
(4) Because responses to experimental drought correspond strongly with ontogeny, seedlings are less drought tolerant than adults, and clinal responses to drought will be more pronounced at the seedling and juvenile stages than for adults.

## Materials and Methods

### Study species

*Conyza canadensis* is an annual, selfing species in the Asteraceae family that primarily colonizes ruderal habitats (Weaver 2001). The native range of *C. canadensis* covers much of North America and its non-native range includes most of the rest of the northern hemisphere. In its native range *C. canadensis* is highly restricted to roadsides, along train tracks, agricultural fields and other highly disturbed areas. In some parts of its non-native range it can be invasive and suppress natives (Shah et al. 2014). Tolerance to abiotic stress, including drought, may be an important mechanism of invasion success of *C. canadensis* (Rosche et al. 2019). A recent precipitation experiment suggested that drought indirectly increases *C. canadensis* abundance by attenuating competitors in its non-native ranges (Mojzes et al. 2020). *Conyza canadensis* is a good model species to study clinal variation as it exhibits high potential for rapid adaptation to a wide range of environmental conditions (Bajwa et al. 2016; Rosche et al. 2019) and biotic interactions (Sheng et al. 2022), possibly because it has a small genome with a large number of genes (Laforest et al. 2020) that may promote rapid evolution (Peng et al. 2014).

### Sample populations

30 native and 29 non-native populations from 17 different geographical regions across the distribution of *C. canadensis* in the northern hemisphere were sampled (Fig. 2a; Suppl. Table 1). Sampling of populations from several distinct geographical regions within either range allows setting population nested within region as a random effect which was recently shown to appropriately account for spatial auto-correlation in *C. canadensis* (Rosche et al. 2019). In each range, populations were separated by a minimum of 10 km and occurred across a broad climatic gradient, including six out of the nine Whittaker biomes (Fig. 2b). From each population, matured seeds from five plants (in total 295 plants) that were selected randomly across an area that was approximately 30 × 30 m in size were sampled. To estimate the aridity patterns experienced by populations (i.e., local aridity regime), monthly climatic water deficit data were sampled for populations using the TerraClimate dataset (Abatzoglou et al. 2018, Young et al. 2021) at a spatial resolution of 1/24° (∼4 km). There was no difference (*p* = 0.989) in the mean annual water deficit between native and non-native populations (Fig. 2c).

**Figure 2.**
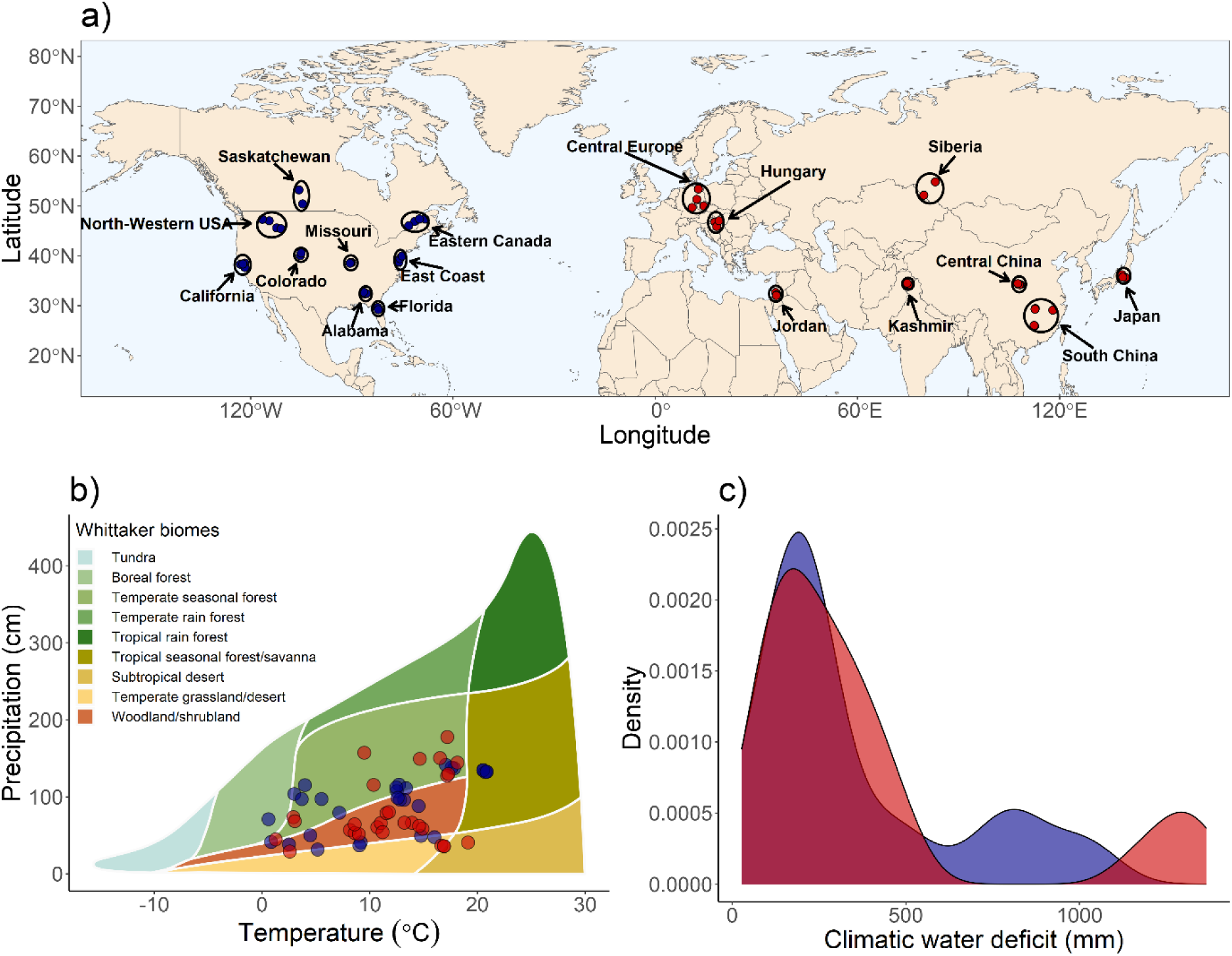
Spatial and bioclimatic distribution of 30 native (blue) and 29 non-native (red) *Conyza canadensis* populations used in the present study. In panel (a), geographical distribution of the populations across nine native and eight non-native regions are presented. In panel (b), populations are plotted according to their mean annual temperature and precipitation on a Whittaker diagram showing the classification of the main terrestrial biomes. In panel (c), the comparison of climatic water deficit between native vs. non-native populations is presented. Climatic water deficit data was downloaded from the TerraClimate database (Abatzoglou et al. 2018), annual precipitation and annual mean temperature data was downloaded from the CliMond database (Kriticos et al. 2012).

### General experimental design

To avoid immediate maternal effects from the field, a potential bias in offspring fitness in common garden experiments (de Villemereuil et al. 2016), F1-offsprings were produced. To do so, seeds from the five seed families per population were germinated in a greenhouse at Martin Luther University Halle-Wittenberg (MLU Halle) in 2020 by covering inflorescences with a mesh bag until achenes were mature to prevent cross-pollination. Of the 295 seed families sampled, 270 produced seeds. These F1-offsprings were used for the experiments.

To compare the effects of experimental drought across life stages, three separate experiments were conducted in parallel in 2021 focusing on (i) seeds and seedlings (recruitment life stage), (ii) juveniles (three weeks after germination) and (iii) adult plants (two months after germination) (Fig. 3). In each experiment, 540 plants were included (270 seed families × 2 experimental drought treatments (wet, dry) using independent seeds from the same seed families. In each experiment, three performance measures and three functional traits were assessed (Fig. 3).

**Figure 3.**
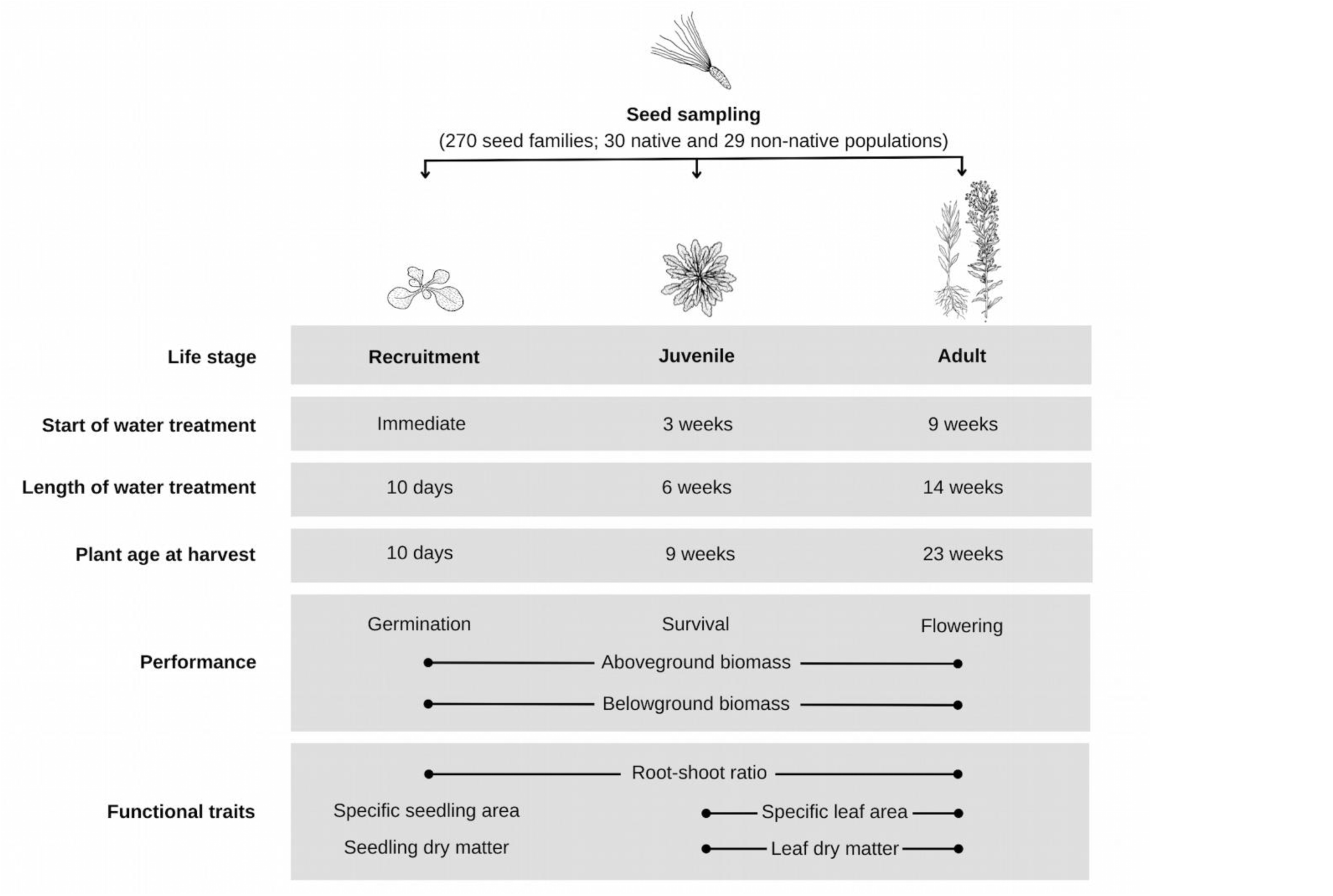
Flowchart illustrating the design of the three experiments and data collection. The drought response of *Conyza canadensis* was tested with two levels using wet vs. dry conditions.

### (i) Development at the recruitment stage

A germination trial was performed to investigate how plant performance and functional traits at the recruitment life stage responded to the wet vs. dry experimental treatments and how these responses correlated with native vs. non-native range and the water deficit of the populations. Ten seeds per seed family were germinated in separate Petri dishes, on filter paper (Whatman No. 1). Two experimental drought treatments were applied using tap water for wet conditions (∼ 0 MPa) and mannitol solution for dry conditions (–0.8 MPa). The dry treatment’s osmotic potential was determined to affect but not inhibit germination and seedling development in a preliminary experiment (Suppl. Fig. 2). Dishes were sealed with Parafilm, re-saturated with tap water daily to maintain consistent concentrations, and kept in a germination chamber at 20 °C at 12 h day and 10 °C at 12 h night. Positions of Petri dishes were randomized daily. Germination (i.e., when radicle breaks through seed coat) was recorded daily with a stereomicroscope. The germination trial ended when no new germination had occurred for five consecutive days.

To assess traits at the recruitment life stage of *C. canadensis*, one seedling per seed family was transferred to another Petri dish and maintained under the same experimental conditions (0 vs. –0.8 MPa). After 10 days, the roots (hypocotyl) were separated from the shoots (epicotyl). Shoots were immediately weighed, and then the shoot area was scanned using WinFolia software (Regent Instruments, Quebec, CA). Roots and shoots were placed separately in paper bags and dried (48h at 60°C) to record dry mass. Three performance traits (germination, above– and belowground biomass) and three functional traits (specific seedling area (SSA), seedling dry matter content (SDMC) and root-shoot ratio (RSR)) were determined (Fig. 3).

### (ii) Growth of juveniles

A greenhouse experiment was performed to investigate how performance and functional traits at the juvenile life stage responded to the wet vs. dry experimental treatments and how these responses correlated with range and water deficit. 540 pots (Stuewe and Sons, Tangent, Oregon, USA; Deepot, D40H, 6.3 × 25 cm, 650 mL) were filled with a water-saturated mixture of sand and local field soil in a 1:1 ratio until a total weight of 600 g. From each seed family, seeds were sown in two pots (2 treatments × 270 seed families = 540 pots). After germination, individuals were randomly thinned out, leaving one individual per pot. Pots were watered equally every two days until the pots were transferred to either the wet or dry treatment which was when the rosette diameter reached an average of 2 cm.

To have comparable conditions at the start of the treatment, all pots for both the wet and dry treatments were again watered until saturation (∼ 30% soil water content measured with Theta Probe Typ ML2x Soil Moisture Device (Delta T Devices, Cambridge UK)). After that saturation, the experimental drought was applied in the following way: Every Monday, Wednesday, and Friday, 59 randomly chosen pots (one from each population) from the wet treatment were weighed. The average pot weight was then subtracted from the average pot weight at saturation (600 g); this difference was the average water loss. This amount of water in mL was added individually to each pot in the wet treatment using a bottle-top dispenser, while the dry treatment pots received 25% of the water added in the wet treatment. During the six-week juvenile experiment, a total of 507 mL water was added to each wet-treatment pot and 126.75 mL water to the dry-treatment pots, roughly equivalent to 213.4 mm vs. and 53.3 mm in precipitation in six weeks. This corresponds to the mean precipitation of the humid South Chinese populations (208.1 mm) and dry Northwestern USA populations (53.4 mm) in May (TerraClimate). The position of single pots within trays and the position of the trays within the greenhouse were randomized weekly.

At the harvest, an average size leaf from each individual was collected, measured for fresh biomass and leaf area with WinFolia software. Roots and shoots were also harvested and, together with leaf samples, dried for 2 days at 60°C, and then weighed. Three performance traits; survival, aboveground biomass, and belowground biomass, and three functional traits; specific leaf area (SLA), leaf dry matter content (LDMC), and root-shoot ratio (RSR) were measured or calculated (Fig. 3).

### (iii) Growth of adults

Another greenhouse experiment was performed to investigate how performance and functional traits at the adult life stage respond to wet vs. dry treatments and how these responses correspond with range and water deficit. 540 pots (1450 mL, Lamprecht-Verpackungen GmbH, Göttingen, Germany, 11 × 11 × 12 cm), were set up in the greenhouse. The pots were filled with a water-saturated mixture of sand and local field soil in a 1:1 ratio, weighing 1300 g. From each seed family, seeds were sown in two pots (2 treatments × 270 seed families = 540 pots). After germination, individuals were thinned out leaving one individual in each pot. Pots were watered equally with a sprayer every two days until they were transitioned to either the wet or dry treatment, which was two months after peak germination, i.e., the same time at which pots with juveniles were harvested.

The experimental drought was started to be applied two months after peak germination, the same time at which juveniles were harvested. Experimental drought and pot randomization were the same as for the juvenile experiment. Through the 14-week adult experiment, a total of 3,652 mL of water was added to each wet pot and 913 mL water to each dry pot, which correlates to a total of 365.2 mm vs. 91.3 mm in precipitation in 14 weeks. This is similar to the warm mesic Alabama populations (335.4 mm) and the dry northwestern USA populations (117.4 mm) sites in May through July.

Plants were harvested 14 weeks after the experimental drought was initiated, at the peak of the flowering. During the harvest, an average size leaf of each individual was collected, measured for fresh biomass and leaf area with WinFolia software. Roots and shoots were also harvested and together with the leaf samples dried for 2 days at 60°C, after which dry mass was recorded. Three performance traits; flowering, aboveground biomass, and belowground biomass, and three functional traits; specific leaf area (SLA), leaf dry matter content (LDMC), and root-shoot ratio (RSR) were measured or calculated (Fig. 3).

### Controlling juvenile and adult experiments

In addition to the experimental pots, 118 additional pots (59 populations × 2 drought treatments) were set up for monitoring soil moisture, every Monday for the juvenile and every second Monday for the adult experiment using a Theta Probe Typ ML2x Soil Moisture Device (Delta T Devices, Cambridge UK) (Suppl. Fig. 3a,b). These 118 pots were treated equally to the experimental pots and were solely used for the purpose of monitoring because the soil moisture measures may cause disturbance which needed to be ruled out as confounding effects on plant performance. Once a week for juveniles and every two weeks for adults, the weight of the 118 experimental pots was recorded, to monitor changes in weight, as another indicator of soil moisture content (Suppl. Fig. 3c,d). At the harvest, stomatal conductance and leaf δ^13^C isotope ratio in every plant individual were measured additionally to evaluate the effects of the wet vs. dry treatments (Suppl. Fig. 3e,f,g,h).

To measure the effect of experimental drought on the experimental plants, the stomatal conductance was measured using an AP4 Porometer (Delta T Devices, Cambridge UK). The largest leaf was considered for the measurement. The porometer was calibrated before each measurement. Carbon stable isotope compositions (δ13C) serves as a proxy for stomatal conductance with water stressed plants having reduced conductance and thus discriminating less the heavier carbon isotope and having higher δ13C values than well-watered plants. Therefore, the δ^13^C isotope ratio was determined from grounded leaf samples with an elemental analyzer Flash 2000 with a TCD detector that was coupled to a Conflo IV and mass spectrometer Delta V Advantage (Thermo Fisher Scientific, Bremen, Germany) at the Center for Stable Isotope Research of Charles University, Prague. The carbon isotope ratio is expressed as follows: Δ^13^C = (R_sample_ / R_standard_ − 1) ∗ 1000, where R is the relative abundance of the carbon isotopes (R = ^13^C/^12^C). The isotope ratio was normalized by using international standards; the carbon isotope ratios were reported using the Vienna Pee Dee Belemnite (VPDB) scale. In addition to the repeated measurements of a series of international standards (e.g., IAEA-CH3, IAEA-CH6 and IAEA-600), a glycine standard was run after every 10th sample to calibrate the elemental composition determinations and to quality control for the isotopic measurements. The analytical precision was within ±0.2‰ for the isotope ratios (Suppl. Fig. 3g,h).

### Statistical analyses

First, it was evaluated whether annual or seasonal water deficit values of the sampled populations were the best indicator of the local aridity regime of the populations. To this end, the models’ performances were compared based on the Akaike information criterion (AIC) with the lowest AIC indicating the best fit. The models tested how the performance and functional traits (six per life stage) were related to the three-way interaction of range × experimental drought × water deficit. For water deficit, five values were used, i.e., the annual water deficit and four seasonal water deficits for three-month-long periods (March-May as spring, June-August as summer, September-November as autumn and December-February as winter). the AIC values for each response were then compared. Overall, the annual water deficit values provided the lowest AIC values across the 18 response variables (Suppl. Table 2.), which supported the use of this estimate in the statistical analyses.

Performance and functional trait analyses were conducted with linear mixed-effects models using the R-package lme4 1.1-29 (Bates et al. 2015). Explanatory variables were the interactive effects of range, experimental drought and centered water deficit. Random factors were population nested within region. Stepwise backward model selection was used based on χ^2^-tests. Normality of residuals and variance homogeneity was evaluated by visually inspecting quantile plots and residual histograms for the models with transformed vs. untransformed variables. Variables remained untransformed if not otherwise stated below. Germination, survival and flowering were analyzed as binomial response variables with generalized linear mixed-effects models. Aboveground biomass and belowground biomass were log-transformed response variables while RSR, SLA (or SSA in case of seedlings) and LDMC (or SDMC in case of seedlings) were non-transformed response variables.

Germination was also analyzed in terms of timing using a time-to-event analysis performed with mixed-effects Cox models with the R package coxme 2.2-16 (Therneau 2012). The same model structure (explanatory variables and random effects) as for the mixed-effects models above was used, again applying stepwise backward model selection based on χ^2^-tests. Cox proportion hazards were checked via Kaplan-Meier plots and multicollinearity was checked via variance inflation factor (VIF) for each model individually (McNair et al. 2012).

In addition to analyzing the recorded absolute values of performance and functional traits, plasticity indices for every measure were calculated to evaluate plasticity in response to experimental drought (Wang & Callaway 2021). The simplified relative distance plasticity index (RDPIs) was chosen because this index is particularly suitable for comparing variation among populations (Valladares et al. 2006). RDPIs for a given trait was calculated as:

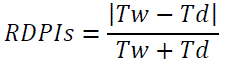

where Tw and Td represented mean trait values for each population in the wet and dry treatments, respectively. The RDPIs-values were then tested for differences between ranges with linear mixed-effects setting region as a random effect.

To investigate whether the effect of the experimental drought on plant performance was determined by functional traits, Pearson correlations between the log response ratios of the measured performance and functional traits to the experimental drought were used. These correlations were recorded separately for each of the three life stages.

To compare the life stages for the effects of the experimental drought and clinal variation, four separate linear mixed-effects models were run for the log-response ratios of each performance measures. The maximal models were the: 1) three way interacting effects of range × centered water deficit × life stage, two way interacting effects of 2) centered water deficit × life stage and 3) range × life stage and the 4) main effect of life stage were tested.

## Results

### Effects of experimental drought on performance and functional traits

Experimental drought affected plant performance in eight out of nine cases (three measurements × three life stages) (Fig. 4, for details of the models see Suppl. Table 3). Across life stages and ranges, plants grown in wet treatments produced 50–450% more biomass than plants in dry treatments. Germination rate was the only trait not affected by experimental drought (Fig. 4a). However, the time-to-event analysis suggested a greater and faster germination through time in wet compared to dry conditions: on average, seeds in the wet treatment germinated one day earlier than seeds in the dry treatment (Suppl. Table 4, Suppl. Fig. 4).

**Figure 4.**
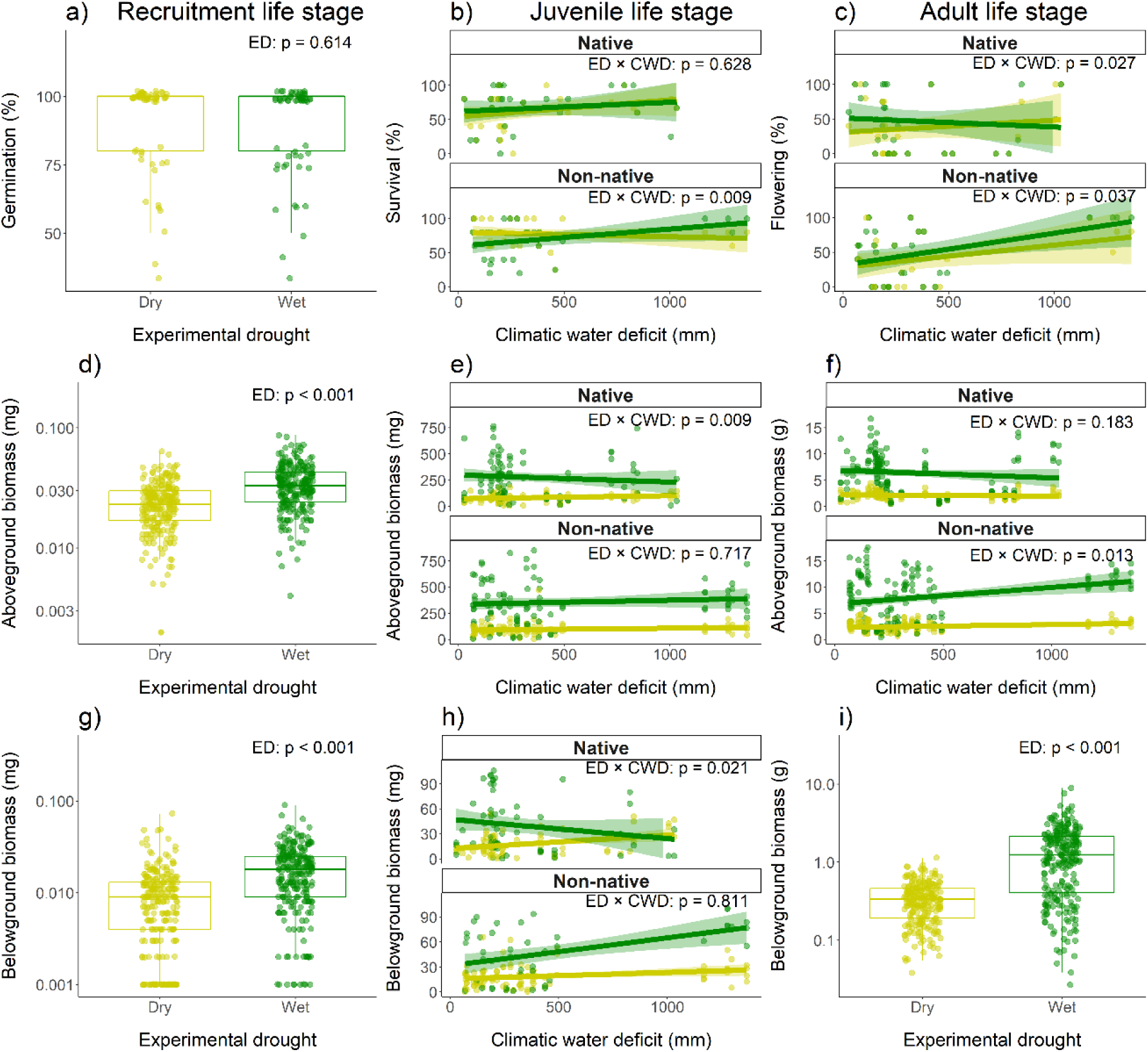
Interactive effects of experimental drought (ED), climatic water deficit (CWD) and range-affiliation (native vs. non-native) on plant performance for germination (a), survival (b) and flowering (c), aboveground biomasses (d-f) and belowground biomasses (g-i). Performance traits were recorded across three experiments focusing on the following life stages: recruitment life stage (a, d, g), juvenile life stage (b, e, h) and adult life stage (c, f, i). The figures represent the results of the minimum adequate models (i.e., including only significant terms), except for the response variables where none of the three tested explanatory factors or their interactions had a significant effect (here: germination rate (a)). For germination, survival and flowering rate, data was analyzed at the individual level (binomial), but population means were plotted as percentages for the sake of clarity. Coloring is based on the applied experimental drought (dry–yellow; wet–green). Details on the models and results can be found in Suppl. Table 3.

Experimental drought affected functional traits in eight out of nine cases (three measurements × three life stages) (Suppl. Table 5, Suppl. Fig. 5). Only LDMC at the adult life stage was not affected by the experimental drought (Suppl. Fig. 5i). There was a shift towards functional traits related to increased growth and resource acquisition by plants in the wet treatment: 14-30% higher SLA, and 22–33% lower LDMC than in the dry-treatment plants (Suppl. Fig. 5g-h). Moreover, plants in the dry treatment had 16–32% higher RSR than plants in the wet treatment (Suppl. Fig. 5a-c).

Correlations between the log-response ratios (i.e., response to the experimental drought) of the performance and the functional traits were significant for only four out of the 27 possible cases (Suppl. Fig. 6). This finding suggests that responses of functional traits to drought did not clearly affect the growth of plants from different populations in wet vs. dry experimental conditions.

### Clinal variation for native and non-native populations

Five out of nine performance measurements (three measurements × three life stages) showed significant three-way interactions among range, experimental drought, and water deficit (Fig. 4, Suppl. Table 3). There was no two-way interaction between water deficit and experimental drought. In other words, whenever clinal variation in the effect of experimental drought on plant performance was present (e.g., Fig. 1c or d), the patterns differed between native and non-native ranges (as in Fig. 1e). In particular, for the native populations, the pattern of clinal variation follows evolutionary expectations from local adaptation for adult flowering, aboveground and belowground biomass of juveniles (Fig. 4c,e,h). In contrast, for non-natives, no such signatures of clinal variation for the mentioned performance measures were found. For non-natives, local clinal variation for juvenile survival, adult flowering and aboveground biomass of adults was present (Fig. 4b,c,f), but this did not follow the evolutionary expectations as ΔP (performance difference between wet and dry treatments) increased with aridity in the population history. For these responses, however, there was no clinal variation in the native populations.

Notably, as main effects, none of the nine performance measures differed between native and non-native populations (e.g., non-natives did not grow bigger). Furthermore, plasticity in response to experimental drought did not differ between native and non-native populations for any of the nine performance measures (Suppl. Fig. 7). This result means there were no mean differences between native vs. non-native populations in performance and their response to experimental drought; yet, how population history affected clinal variation differed strongly across the ranges.

Functional traits did not show a consistent pattern of clinal variation across life stages. Only SLA of adults (Suppl. Fig. 5f) produced a significant three-way interaction, in which the non-native plants showed clinal variation, but the native plants unexpectedly did not. Seedling dry matter content was the only trait with a significant two-way interaction between experimental drought and water deficit (Suppl. Figure. 5g). This result suggests similarly pronounced clinal variation between the ranges for this functional trait. With range as a main effect, none of the nine functional traits differed between native and non-native populations, nor did plasticity in response to experimental drought differ between native and non-native populations for any of the nine functional traits (Suppl. Fig. 8).

**Figure 5.**
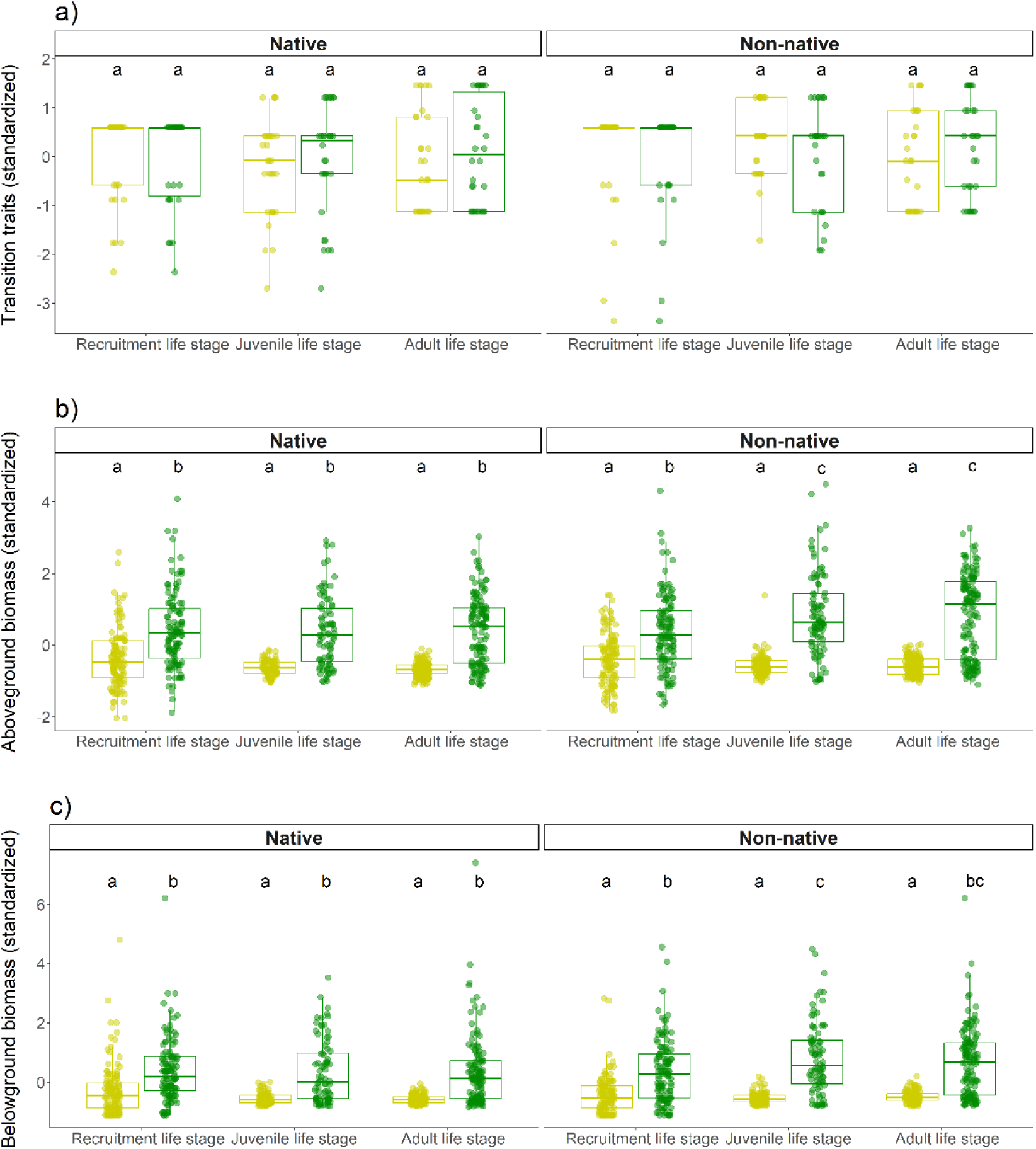
Life stage (recruitment, juvenile and adult) and their interactive effects with range and drought treatment on performance, such as the binomial transition traits (germination, survival and flowering (a)), aboveground-(b) and belowground (c) biomasses. The models tested the differences in log response ratios, however, the plot aimed to visualize the comparable effect of experimental drought across life stages. To reduce the effect of scale differences among life-stages (e.g., biomass variation), all variables were standardized across life-stages. For transitional traits, binomial data was analyzed but population means were plotted for the sake of clarity. Coloring is based on the applied experimental drought (dry–yellow; wet–green). Details on the models and results can be found in Supplementary Table 6. Lowercase letters refer to the groupings tested with posthoc analyses (Tukey HSD) performed on the standardized data.

### Life stages and clinal variation

For each of the three performance traits, the effect of experimental drought (i.e., log-response ratios between wet vs. dry treatments) differed among the life stages (Suppl. Table 6). There were overall smaller log-response ratios for the recruitment life stage than for the juvenile life stage (by 48.1–95.8%) and the adult life stage (by 41.7 – 95.5%). However, when comparing individual traits between wet and dry treated plants, consistent patterns across all life stages were found. In particular, germination, survival, flowering did not differ between treatments. In contrast, aboveground and belowground biomass was 50-450% greater in wet than in dry treatment across all life stages (Fig. 5). More importantly, log-response ratios were not affected by an interaction between life stage and water deficit or life stage and range or their three-way interactions (Suppl. Table 6). These results indicate that the patterns of clinal variation were similar among the life stages.

## Discussion

This research is the first to investigate clinal variation in response to experimental drought across several life stages while studying broad biogeographical scales in both native and non-native ranges. Drought reduced performance consistently at all life stages and across both ranges. Patterns of clinal variation were consistent across the life stages but contrasting patterns of clinal variation between the two ranges were found. Native populations from xeric habitats were less inhibited by drought than mesic populations, displaying the expected clinal variation. In contrast, non-native populations did not follow this pattern.

## Effects of experimental drought on performance and functional traits

Experimental drought had consistent strong effects on plant performance across all life stages, setting the stage for testing the four predictions. In terms of functional traits, experimental drought decreased SLA and increased LDMC which is a common response of plants to increase stress tolerance by reducing fast growth (Balachowski & Volaire 2018). Root-shoot ratio was reduced by drought, which was consistent across life stages but is not consistent with other studies (Knight et al. 2006; Qi et al. 2019). However, investing resources in increasing RSR under drought may be more advantageous for perennial plants than for short-lived annuals, as the developed root system may be more cost-effective in multiple years than in a single season (Mokany et al. 2006). Instead, *C. canadensis* may respond by more quickly completing its annual life cycle (Brandenburger et al. 2022). Such response would suggest a drought avoidance strategy, which often plays an important role in drought survival (Volaire 2018).

The weak relationships between functional trait response and performance suggest that the measured functional traits have either weak or complex effects on how *C. canadensis* copes with drought stress. Drought stress may require multidimensional responses that may not be captured by single functional traits (Kooyers 2015). Moreover, different functional traits may be subject to contrasting climate selection pressures, and trait correlations and ecological trade-offs may prevent functional traits from adapting linearly to distinct climatic gradients (Ahrens et al. 2020). Similar to the research of this present study, a recent study found patterns of clinal variation in arbuscular mycorrhizal fungi (AMF) colonization in *C. canadensis* but these patterns could not be linked to their measured plant functional traits (Sheng et al. 2022).

## Performance and plasticity in native vs. non-native ranges

There were no general differences in common garden performance between native vs. non-native populations, which contrasts with many studies of other invasive species (reviewed by Parker et al. 2013; Callaway et al. 2022), and a study that found the opposite for *C. canadensis* (Abhilasha & Joshi 2009). However, differences between populations from native vs. non-native ranges may not be present when very large and comparable environmental gradients in both distribution ranges are considered, as recently demonstrated for *C. canadensis* (Rosche et al. 2019, Sheng et al. 2022). Also, benign greenhouse conditions might mask differences in growth between populations from different ranges.

Plasticity of responses to the experimental drought was comparable between the two ranges. This result does not match the assumption that increased plasticity of non-native populations is an important factor for invasion success in general (Richards et al. 2006) and in response to climatic conditions (Turner et al. 2015). Apparently, high-performance genotypes or greater plasticity in non-native populations are not the primary reason for the overwhelming colonization success of *C. canadensis* in the Northern hemisphere. Instead, shifts in biotic interactions such as competition (Shah et al. 2014; Nagy et al. 2022) and/or plant-soil feedback (Sheng et al. 2022) are more likely to drive the success of non-native *C. canadensis* populations.

## Clinal variation in native vs. non-native ranges

Five out of nine performance measures showed clinal variation in response to experimental drought. Although this study did not explicitly test for local adaptation, the results indicate that *C. canadensis* shows large adaptive variation to climate, which may have facilitated its ability to occur across the unusually large climatic gradient of its cosmopolitan distribution (see also Rosche et al. 2019). In all five cases, this clinal variation differed between the two ranges. The response of native populations to experimental drought met adaptive and evolutionary expectations in the native range – populations from xeric habitats were less suppressed by the dry treatment. Such clinal variation was not apparent in non-native populations. These results are consistent with studies of *C. stoebe* (Mráz et al. 2014) and *P. lanceolata* (Villellas et al. 2021), suggesting generality in range-specific clinal variation.

This study postulates two mutually non-exclusive mechanisms that may explain why non-native populations may not have locally adapted, or perhaps yet, to variation in water supply. First, non-native populations have been introduced to their current climatic regimes much more recently (Yan et al. 2020), and that did not necessarily match their previous adaption to climate. Thus, non-native populations may have not experienced enough time to adapt to their new abiotic environments. Second, the relative importance of different selection regimes may differ between ranges. For example, fundamentally altered biotic interactions – such as a lack of specialized enemies in the non-native ranges – may be more important for the non-native *C. canadensis* populations than a “perfect adaptation” to local aridity (see also Callaway et al. 2011, Pal et al. 2020). Such basic evolutionary research questions need to be addressed with more complex experimental designs that simultaneously manipulate abiotic and biotic conditions. For example, studying competitive interactions simultaneously with responses to drought for many native vs. non-native populations across large geographic distributions could enhance our mechanistic understanding of rapid evolution.

Irrespective of these mechanisms, these results have two important implications. First, *C. canadensis* is a highly successful weed in its non-native distribution. Assuming that it is not yet fully adapted to climate, the spread and impact of this species might become even more pronounced in the future. Second, *C. canadensis* has occurred in some non-native ranges for several centuries and is actually known to be able to rapidly respond to certain environmental changes (Rosche et al. 2019). If even this species has not yet established clines related to drought in the non-native ranges, it remains questionable if and how many plant species can evolve fast enough to deal adequately with climate change which is also occurring at contemporary time scales. Such interpretation is concerning with a view on global change scenarios and calls for further investigations in comparable future studies.

## Drought response and clinal variation across life stages

Experimental drought had weaker effects in the recruitment life stage than in the other two life stages. Plants may respond less plastically at the recruitment stage where they must either grow very rapidly or die as found for *Solidago gigantea* (Nagy et al. 2018). In contrast, above-and belowground biomass was reduced, and transitional traits not affected for all life stages. Range-specific clinal variation was also similar across life stages. This study predicted more variation among life stages, as different life stages appear to need different strategies to tolerate water stress (Coe et al. 2021), and adaptive differentiation should be particularly strong during recruitment when young plants are vulnerable to abiotic stress (Poorter 2009, Postma and Ågren 2016). However, similarly pronounced drought responses and clinal variation across life stages could be particularly important under climate change because it is expected to lead to more extreme weather events, such as unpredictable droughts at any life stage (Parmesan & Hanley 2015). If a plant species responds similarly to these different weather conditions across all life stages, it will be better able to adapt and survive (Garzón et al. 2011; Moran et al. 2016; Welles & Funk 2021). This may have several positive effects for the colonization of large ranges and adaptive strategies to the resilience of species to changing climatic conditions.

## Conclusions

The results of this study shed light on the ecological and evolutionary mechanisms underlying the adaptation of invasive species in new environments, and the speed of adaptation to drought. The experiments suggest that invaders can thrive even in the absence of complete adaptation to new abiotic environments, indicating their remarkable resilience in the face of changing global conditions. Furthermore, our findings suggest that long-established invaders may continue to evolve in response to the dynamic abiotic environment. Future research is needed to test the generality of these findings with focus on 1) the implications for the adaptive potential of plant populations under ongoing climate change, 2) whether invasion success of some non-native species become even more pronounced once they are fully adapted to the local climate in the novel ranges, and 3) the relative importance of abiotic and biotic selection drivers for rapid evolution in non-native plants.

## Acknowledgements

The project was supported by the Deutsche Forschungsgemeinschaft (DFG, German Research Foundation; project number: RO 6418/1-1 to C.R.), an MLU|BioDivFund grant (R02020830-29 to C.R.), an NSERC grant (RGPIN-2022-05262 to D.P.K). M.A.S. acknowledges the support through RUSA 2.0 project by Mission Directorate RUSA J&K and Ministry of Education, Govt. of India. I. H and C.R. gratefully acknowledge the support of iDiv funded by DFG (FZT 118, 202548816). We also wish to thank to Rachel Draude, Heidi Hirsch, Katrin Kittlaus, and Anne Sophie Winkler for their help during the experiments and manuscript visualization.

## Author contribution

DUN and CR designed and planned the experiment. DUN, MAG, SLF, LJF, MH, DPK, YL, RWP, MAS, MSH, MSL, CS, TS and CR conducted field work and sampling. DUN, MH and CR analyzed the data. DUN, AET, KJ, IS and CR performed the experiments. DUN, RMC, SLF, IH, YL and CR wrote the manuscript.

## Notes

### Competing Interest Statement

The authors have declared no competing interest.

