## Supporting information for "Clinal variation in drought response is consistent across life stages but not between native and non-native ranges"

**Article acceptance date:** Not yet accepted

**The following Supporting Information is available for this article:**

**Supplementary Table 1.** Geographical location and climatic water deficit of 30 native and 29 non-native *Conyza canadensis* populations.

**Supplementary Table 2.** Model selection by Akaike information criterion values testing how seasonal and annual climatic water deficit influence the models.

**Supplementary Table 3.** Details of the linear mixed-effects models on performance measures.

**Supplementary Table 4.** Details of the linear mixed-effects Cox model.

**Supplementary Table 5.** Details of the linear mixed-effects models on functional traits.

**Supplementary Table 6.** Details of the linear mixed-effects models on log-response ratios.

**Supplementary Figure 1.** Concept of clinal variation in the experiment.

**Supplementary Figure 2.** Results of germination and early seedling traits in a pre-experiment.

**Supplementary Figure 3.** Measurements of the effects of experimental drought, during the juvenile and adult life stages.

**Supplementary Figure 4.** Kaplan-Meier plot for germination success.

**Supplementary Figure 5.** Interactive effects of experimental drought, climatic water deficit and range-affiliation on plant functional traits.

**Supplementary Figure 6.** Details of the analyses investigating the correlation among performance and functional traits.

**Supplementary Figure 7.** Effects of range-affiliation on plasticity of plant performance.

**Supplementary Figure 8.** Effects of range-affiliation on plasticity of functional traits.

**Supplementary Table 1.** Geographical location and climatic water deficit (CWD) of 30 native and 29 non-native *Conyza canadensis* populations used in the present study. CWD was summed to detect seasonal differences for Spring (March-May), Summer (June-August), Autumn (September-October), Winter (December-February) and annual (the whole year). Native populations are assigned to nine geographic regions: California (CA), North-Western USA (NU), Saskatchewan (SK), Colorado (CO), Missouri (MO), Alabama (AL), Florida (FL), East Coast (EC) and Eastern Canada (NB and QU). Non-native populations are assigned to eight geographic regions: Central Europe (CE), Hungary (HU), Jordan (JO), Kashmir (KA), Siberia (SB), Central China (CC), South China (SC) and Japan (JP).

| Pop ID | Region | Location | Latitude | Longitude | CWD (Annual) | CWD (Spring) | CWD (Summer) | CWD (Autumn) | CWD (Winter) |
|---|---|---|---|---|---|---|---|---|---|
| *Native range* | | | | | | | | | |
| AL-HL | Alabama | Montgomery | 32.37 | -86.15 | 187.43 | 0.23 | 37.19 | 89.78 | 0.96 |
| AL-PR | Alabama | Tuskegee | 32.47 | -85.69 | 165.78 | 0.19 | 33.53 | 78.47 | 0.80 |
| AL-VE | Alabama | Verbena | 32.74 | -86.53 | 152.87 | 0.19 | 29.59 | 78.28 | 0.75 |
| CA-BB | California | Bodega Bay | 38.31 | -123.06 | 416.64 | 2.30 | 66.09 | 228.83 | 7.14 |
| CA-DA | California | Davis | 38.54 | -121.78 | 1032.44 | 3.38 | 189.77 | 600.36 | 9.84 |
| CA-LV | California | Livermore | 37.66 | -121.73 | 1006.2 | 5.04 | 196.47 | 562.75 | 14.15 |
| CA-NV | California | Napa Valley | 38.37 | -122.34 | 845.73 | 2.70 | 122.78 | 514.69 | 8.07 |
| CO-BO | Colorado | Boulder | 40.03 | -105.29 | 785.52 | 9.18 | 152.11 | 423.88 | 21.29 |
| CO-FC | Colorado | Fort Collins | 40.56 | -105.04 | 827.61 | 1.76 | 185.12 | 429.33 | 10.12 |
| EC-EA | East Coast | Easton | 38.75 | -76.07 | 196.36 | 0.86 | 35.78 | 116.68 | 1.85 |
| EC-MT | East Coast | Middletown | 39.44 | -75.74 | 187.42 | 0.29 | 32.90 | 116.41 | 0.98 |
| EC-PH | East Coast | Philadelphia | 39.97 | -75.12 | 157.76 | 0.15 | 29.07 | 98.42 | 0.56 |
| FL-AN | Florida | Anthony | 29.26 | -82.06 | 200.17 | 10.82 | 111.78 | 14.98 | 25.97 |
| FL-BA | Florida | Bivens Arm | 29.63 | -82.35 | 193.59 | 8.43 | 115.47 | 12.23 | 20.15 |
| FL-CT | Florida | Citra | 29.41 | -82.15 | 197.67 | 9.95 | 113.93 | 13.69 | 23.35 |
| FL-GV | Florida | Gainesville | 29.62 | -82.34 | 188.5 | 8.39 | 113.38 | 11.82 | 19.46 |
| FL-OC | Florida | Ocala | 29.21 | -82.15 | 208.24 | 11.34 | 117.44 | 13.90 | 27.46 |
| MO-FP | Missouri | Fairmont City | 38.66 | -90.08 | 218.46 | 0.22 | 27.40 | 132.29 | 0.51 |
| MO-SL | Missouri | St. Louis | 38.68 | -90.36 | 213.38 | 0.12 | 26.64 | 128.93 | 0.46 |
| MO-TY | Missouri | Eureka Tyson | 38.53 | -90.56 | 239.59 | 0.17 | 28.01 | 150.72 | 0.68 |
| NU-AT | North-Western USA | Alberton | 47.02 | -114.51 | 519.02 | 0.00 | 61.92 | 356.09 | 0.00 |
| NU-BT | North-Western USA | Butte | 45.59 | -112.32 | 723.65 | 0.03 | 164.89 | 403.40 | 1.33 |
| NU-BZ | North-Western USA | Bozeman | 45.42 | -111.03 | 242.39 | 0.00 | 0.00 | 177.01 | 0.00 |
| NU-SM | North-Western USA | Saint-Maries, Idaho | 47.31 | -116.56 | 306.47 | 0.00 | 31.99 | 214.49 | 0.00 |
| SK-PA | Saskatchewan | Prince Albert | 53.22 | -105.71 | 258.2 | 0.00 | 41.20 | 169.72 | 0.00 |
| SK-RE | Saskatchewan | Regina | 50.42 | -104.62 | 480.71 | 0.00 | 80.95 | 298.91 | 0.00 |
| NB-ED | Eastern Canada | Edmundston | 47.38 | -68.3 | 56.76 | 0.00 | 2.53 | 50.36 | 0.00 |

| | | | | | | | | | |
|---|---|---|---|---|---|---|---|---|---|
| QU-LB | Eastern Canada | Lac Beauport | 46.91 | -71.32 | 26.97 | 0.00 | 2.98 | 23.73 | 0.00 |
| QU-LP | Eastern Canada | La Pocatière | 47.38 | -69.97 | 83.88 | 0.00 | 4.45 | 74.28 | 0.00 |
| QU-ST | Eastern Canada | Sorel-Tracy | 46.04 | -73.13 | 105.75 | 0.00 | 12.26 | 81.99 | 0.00 |
| *Non-native range* | | | | | | | | | |
| CE-FO | Central Europe | Faule Ort | 53.4 | 12.8 | 151.27 | 0.00 | 28.75 | 103.31 | 0.00 |
| CE-PG | Central Europe | Prague | 50 | 14.41 | 215.65 | 0.24 | 52.03 | 124.83 | 0.72 |
| CE-PW | Central Europe | Plagwitz | 51.33 | 12.32 | 214.32 | 0.31 | 46.67 | 134.24 | 1.04 |
| CE-ZD | Central Europe | Zeckendorf | 49.69 | 10.94 | 137.29 | 0.00 | 25.69 | 95.19 | 0.00 |
| HU-BA | Hungary | Balatonlelle | 46.79 | 17.71 | 290.2 | 0.23 | 63.69 | 184.05 | 0.94 |
| HU-MA | Hungary | Máriagyűd | 45.88 | 18.25 | 257.75 | 0.17 | 49.33 | 163.80 | 0.69 |
| HU-RA | Hungary | Rácalmás | 47.03 | 18.92 | 379.43 | 0.23 | 82.42 | 237.58 | 0.65 |
| JO-IR | Jordan | Irbid | 32.48 | 35.98 | 1276.8 | 11.54 | 319.23 | 619.84 | 21.92 |
| JO-MK | Jordan | Malka | 32.65 | 35.72 | 1167.95 | 6.95 | 258.91 | 595.24 | 12.48 |
| JO-SA | Jordan | Sarot | 32.16 | 36 | 1365.84 | 19.71 | 363.19 | 631.92 | 40.68 |
| JO-SB | Jordan | Shaffa Badran | 32.06 | 35.93 | 1296.77 | 14.23 | 332.74 | 621.28 | 25.82 |
| JP-NA | Japan | Nagano | 36.22 | 138.48 | 71.07 | 2.14 | 29.50 | 30.12 | 2.86 |
| JP-TK | Japan | Tokyo | 35.65 | 139.46 | 68.25 | 5.77 | 18.15 | 20.69 | 23.21 |
| JP-YA | Japan | Yamanashi | 35.74 | 138.71 | 76.13 | 6.60 | 29.36 | 25.59 | 13.01 |
| KA-GB | Kashmir | Gandarbal | 34.22 | 74.77 | 457.8 | 8.12 | 40.92 | 272.29 | 12.12 |
| KA-HZ | Kashmir | Nigeen Hazratbal | 34.12 | 74.84 | 436.51 | 8.16 | 38.37 | 253.69 | 11.70 |
| KA-SM | Kashmir | Rezan Sonamarg | 34.27 | 75.15 | 244.29 | 0.00 | 9.77 | 149.58 | 0.00 |
| KA-SP | Kashmir | Sopore | 34.54 | 74.73 | 277.84 | 0.00 | 9.76 | 168.86 | 0.00 |
| CC-ME | Central China | Meixian | 34.31 | 107.73 | 320.63 | 9.29 | 119.62 | 145.58 | 28.76 |
| CC-SH | Central China | Shuidaokou | 34.24 | 108.4 | 383.19 | 10.74 | 137.53 | 178.81 | 31.68 |
| CC-YA | Central China | Yangling | 34.24 | 108.05 | 357.32 | 10.28 | 133.18 | 165.53 | 29.66 |
| CC-ZH | Central China | Zhougongmiao | 34.45 | 107.6 | 326.23 | 8.99 | 123.97 | 146.63 | 28.15 |
| SB-AC | Siberia | Aczegul' | 52.16 | 79.71 | 492.57 | 0.00 | 88.75 | 319.93 | 0.00 |
| SB-NS | Siberia | Novosibirsk | 54.84 | 83.1 | 193.17 | 0.00 | 26.84 | 140.39 | 0.00 |
| SC-AO | South China | Aoquanzhen | 25.94 | 112.56 | 121.48 | 8.31 | 0.94 | 44.38 | 14.05 |
| SC-CJ | South China | Cui Jiang Cun | 26.05 | 112.43 | 109.84 | 7.55 | 0.60 | 40.81 | 12.13 |
| SC-QI | South China | Qianlianghuzhen | 29.45 | 112.76 | 151.79 | 7.32 | 3.67 | 70.93 | 13.80 |
| SC-XS | South China | Xingangshan | 29.12 | 117.95 | 156.62 | 9.17 | 1.21 | 72.30 | 12.53 |
| SC-ZH | South China | Zhuzikouzhen | 29.3 | 112.69 | 147.04 | 7.28 | 3.21 | 67.71 | 13.50 |

**Supplementary Table 2.** Model selection by Akaike information criterion (AIC) values testing how seasonal and annual climatic water deficit influence the models. The same model (three-way interaction of range × experimental drought × centered water deficit) was run for all tested response variables at all life-stages, with testing if the application annual, Spring, Summer, Autumn or Winter values result in the best fitting model. After running the models, the AIC values were extracted for each model on the same response variable received ranking scores (1 to 5) to summarize their fit. The lowest AIC got ranking score 1 (dark green), the second lowest 2 (light green), the third lowest 3 (creamy yellow) the second highest 4 (light orange) and the highest value got 5 (dark orange). The ranking scores were then summed up separately for each life-stage and, then for all of them. Overall the annual climatic water balance resulted in the lowest cumulative ranking scores, which suggested using it in the final models. Abbreviations: RSR: root – shoot ratio; SLA: specific leaf area; LDMC: leaf dry matter content; SSA: specific seedling area; SDMC: seedling dry matter content.

| **Adult life stage** | **Annual deficit** AIC | **Spring deficit** AIC | **Summer deficit** AIC | **Autumn deficit** AIC | **Winter deficit** AIC |
|---|---|---|---|---|---|
| Flowering | 507.03 | 509.49 | 507.84 | 507.47 | 516.575 |
| Aboveground biomass | 2387.347 | 2384.927 | 2395.633 | 2380.895 | 2389.029 |
| Belowground biomass | 1174.669 | 1183.749 | 1192.163 | 1184.149 | 1155.831 |
| Root-shoot ratio | -1098.328 | -1098.271 | -1096.569 | -1101.124 | -1099.59 |
| SLA | 3465.59 | 3467.56 | 3469.913 | 3468.929 | 3450.712 |
| LDMC | -1500.512 | -1501.367 | -1499.467 | -1501.228 | -1514.679 |
| Subsummed score | 13 | 19 | 28 | 16 | 14 |
| **Juvenile life stage** | | | | | |
| Survival | 646.516 | 647.61 | 646.894 | 646.5265 | 653.256 |
| Aboveground biomass | -306.2959 | -304.3725 | -303.8538 | -303.9997 | -317.698 |
| Belowground biomass | -975.501 | -962.095 | -965.0727 | -963.0242 | -962.4122 |
| Root-shoot ratio | -219.6848 | -219.1189 | -220.6338 | -220.5244 | -221.222 |
| SLA | 2782.212 | 2783.755 | 2783.44 | 2783.62 | 2780.607 |
| LDMC | -523.6217 | -525.9839 | -524.3412 | -523.2867 | -523.24 |
| Subsummed score | 14 | 25 | 16 | 20 | 15 |
| **Early seedling life stage** | | | | | |
| Germination (binomial) | 313.661 | 313.564 | 313.758 | 313.752 | 309.156 |
| Aboveground biomass | -9093.477 | -9089.494 | -9091.85 | -9091.76 | -9089.022 |
| Belowground biomass | -9147.265 | -9146.72 | -9146.532 | -9149.113 | -9146.02 |
| Root-shoot ratio | 474.1339 | 472.6187 | 474.7116 | 475.0019 | 477.604 |
| SSA | 4134.567 | 4139.15 | 4132.392 | 4135.245 | 4140.491 |
| SDMC | -1562.057 | -1561.474 | -1561.128 | -1561.512 | -1558.94 |
| Subsummed score | 11 | 17 | 19 | 17 | 26 |
| Summed score | 38 | 61 | 63 | 53 | 55 |

**Supplementary Table 3.** Details of the linear mixed-effects models investigating how germination, survival, flowering (binomial), above- and belowground biomass were related to range (native v. non-native), treatment (wet vs. dry), climatic water deficit (CWD) and their interactions, across 30 native and 29 non-native *C. canadensis* populations.

| | Recruitment stage | | | | | | Juvenile stage | | | | | | Adult stage | | | | | |
|---|---|---|---|---|---|---|---|---|---|---|---|---|---|---|---|---|---|---|
| Fixed effects | Germination (binomial) | | $log_e$-transformed aboveground biomass | | $log_e$-transformed belowground biomass | | Survival (binomial) | | $log_e$-transformed aboveground biomass | | $log_e$-transformed belowground biomass | | Flowering (binomial) | | $log_e$-transformed aboveground biomass | | $log_e$-transformed belowground biomass | |
| | $\chi^2_{(1)}$ | Estimate | $\chi^2_{(1)}$ | Estimate | $\chi^2_{(1)}$ | Estimate | $\chi^2_{(1)}$ | Estimate | $\chi^2_{(1)}$ | Estimate | $\chi^2_{(1)}$ | Estimate | $\chi^2_{(1)}$ | Estimate | $\chi^2_{(1)}$ | Estimate | $\chi^2_{(1)}$ | Estimate |
| (Intercept) | | 3.55 | | -10.74 | | -11.93 | | 0.67 | | -2.77 | | -4.58 | | -1.64 | | 0.62 | | -1.34 |
| $Range_{Non-native}$ : $Treatment_{Dry}$ : CWD | 0.02 | n.s. | 2.95 | n.s. | 1.49 | n.s. | 4.17 | 0.95 * | 6.55 | 0.42 * | 3.95 | 0.47 * | 9.07 | 1.76 ** | 6.23 | 0.18 * | 0.94 | n.s. |
| $Range_{Non-native}$ : $Treatment_{Wet}$ | 0.06 | n.s. | 0.05 | n.s. | 0.12 | n.s. | i. int. | -0.58 | i. int. | 0.14 | i. int. | 0.33 | i. int. | -0.06 | i. int. | 0.06 | 0.02 | n.s. |
| $Range_{Non-native}$ : CWD | 0.03 | n.s. | 0.37 | n.s. | 1.21 | n.s. | i. int. | -0.51 | i. int. | -0.15 | i. int. | -0.22 | i. int. | 0.26 | i. int. | 0.09 | 2.28 | n.s. |
| $Treatment_{Wet}$ : CWD | 0.11 | n.s. | 0.51 | n.s. | 0.02 | n.s. | i. int. | -0.17 | i. int. | -0.39 | i. int. | -0.5 | i. int. | -0.97 | i. int. | -0.08 | 1.78 | n.s. |
| $Range_{Non-native}$ | 0.22 | n.s. | 0.01 | n.s. | 0.78 | n.s. | i. int. | 0.8 | i. int. | 0.19 | i. int. | 0.19 | i. int. | 0.59 | i. int. | 0.14 | 0.68 | n.s. |
| $Treatment_{Wet}$ | 0.25 | n.s. | 69.44 | 0.38 | 66.96 | 0.76 *** | i. int. | 0.13 | i. int. | 1.03 | i. int. | 1.19 | i. int. | 0.77 | i. int. | 0.94 | 318.49 | 1.13 |
| CWD | 0.02 | n.s. | 0.4 | n.s. | 5.98 | -0.12 * | i. int. | 0.3 | i. int. | 0.19 | i. int. | 0.38 | i. int. | -0.11 | i. int. | 0.01 | 2.8 | n.s. |
| Random effect variances | | | | | | | | | | | | | | | | | | |
| Region:Population | 3.44 (59) | | 0.02 (59) | | 0.01 (59) | | 0.54 (59) | | 0.07 (58) | | 0.12 (59) | | 0.69 (59) | | 0.01 (59) | | 0.01 (59) | |
| Region | 0.62 (17) | | <0.01 (17) | | <0.01 (17) | | 0.2 (17) | | 0.21 (17) | | 0.47 (17) | | 4.59 (17) | | 0.32 (17) | | 0.65 (17) | |
| Residuals | n.e. (540) | | 0.22 (471) | | 0.93 (470) | | n.e. (540) | | 0.48 (356) | | 0.99 (339) | | n.e. (537) | | 0.15 (528) | | 0.15 (489) | |

Notes: The first column of the table shows the model structure of the maximal models. $\chi2$ -values, parameter estimates, and significance levels of fixed effect terms that remained in the minimal adequate models are given. Range and treatment are denoted with the second factor level. Random effect variances of the region and population nested within region are presented with their number of groups in parentheses (i.e., 59 populations, 17 geographical regions). Random effect variances of residuals are presented with the number of observations. Abbreviations: n.s., not significant; i. int., not estimated because the main factor was in a significant interaction. Significance levels: * $p < 0.05$; ** $p < 0.01$; *** $p < 0.001$.

**Supplementary Table 4.** Details of the linear mixed-effects Cox model investigating how germination dynamics was related to range (native v. non-native), treatment (wet vs. dry), climatic water deficit (CWD) and their interactions, across 30 native and 29 non-native *C. canadensis* populations.

| Fixed effects | Germination dynamics | |
|---|---|---|
| | $\chi^2_{(1)}$ | Estimate |
| (Intercept) | | n.e. |
| $Range_{Non\text{-}native}$ : $Treatment_{Dry}$ : CWD | 0.04 | n.s. |
| $Range_{Non\text{-}native}$ : $Treatment_{Wet}$ | 0.29 | n.s. |
| $Range_{Non\text{-}native}$ : CWD | 0.08 | n.s. |
| $Treatment_{Wet}$ : CWD | 1.26 | n.s. |
| $Range_{Non\text{-}native}$ | 2.85 | n.s. |
| $Treatment_{Wet}$ | 348.58 | 0.74 *** |
| CWD | 0.11 | n.s. |
| Random effect variances | | |
| Region:Population | 0.07 (59) | |
| Region | 0.01 (17) | |
| Residuals | n.e. (4271) | |

Notes: The first column of the table shows the model structure of the maximal models. χ 2 -values, parameter estimates, and significance levels of fixed effect terms that remained in the minimal adequate models are given. Range and treatment are denoted with the second factor level. Random effect variances of the region and population nested within region are presented with their number of groups in parentheses (i.e., 59 populations, 17 geographical regions). Random effect variances of residuals are presented with the number of observations. n.s., not significant; i. int., not estimated because the main factor was in a significant interaction; significance levels: * p < 0.05; ** p < 0.01; *** p < 0.001.

**Supplementary Table 5.** Details of the linear mixed-effects models investigating how root-shoot ratio, specific leaf area and leaf dry matter content were related to range (native v. non-native), treatment (wet vs. dry), climatic water deficit (CWD) and their interactions, across 30 native and 29 non-native *C. canadensis* populations.

| | Recruitment stage | | | | | | Juvenile stage | | | | | | Adult stage | | | | | |
|---|---|---|---|---|---|---|---|---|---|---|---|---|---|---|---|---|---|---|
| Fixed effects | Root-shoot ratio | | Specific seedling area | | Seedling dry matter content | | Root-shoot ratio | | Specific leaf area | | Leaf dry matter content | | Root-shoot ratio | | Specific leaf area | | Leaf dry matter content | |
| | $\chi^2_{(1)}$ | Estimate | | Estimate | $\chi^2_{(1)}$ | Estimate | $\chi^2_{(1)}$ | Estimate | $\chi^2_{(1)}$ | Estimate | $\chi^2_{(1)}$ | Estimate | $\chi^2_{(1)}$ | Estimate | $\chi^2_{(1)}$ | Estimate | $\chi^2_{(1)}$ | Estimate |
| (Intercept) | | 0.44 | | 81.83 | | 11.02 | | 0.19 | | 32.82 | | 26.81 | | 0.15 | | 29.57 | | 18.93 |
| $Range_{Non\text{-}native}$ : $Treatment_{Wet}$ : CWD | 0.65 | n.s. | 2.95 | n.s. | 0.04 | n.s. | 0.39 | n.s. | 0.13 | n.s. | 0.25 | n.s. | 0.64 | n.s. | 5.71 | -0.19 * | 0.06 | n.s. |
| $Range_{Non\text{-}native}$ : $Treatment_{Wet}$ | 0.42 | n.s. | 0.53 | n.s. | 0.37 | n.s. | 0.98 | n.s. | 0.75 | n.s. | 0.22 | n.s. | 1.50 | n.s. | i. int. | 2.12 | 0.28 | n.s. |
| $Range_{Non\text{-}native}$ : CWD | 1.93 | n.s. | 0.26 | n.s. | 0.01 | n.s. | 0.28 | n.s. | 0.02 | n.s. | 1.05 | n.s. | 0.29 | n.s. | i. int. | 3.65 | 1.14 | n.s. |
| $Treatment_{Wet}$ : CWD | 0.49 | n.s. | 1.36 | n.s. | 6.07 | 0.87 * | 0.40 | n.s. | 0.71 | n.s. | 0.90 | n.s. | 2.17 | n.s. | i. int. | 1.87 | 0.03 | n.s. |
| $Range_{Non\text{-}native}$ | 0.22 | n.s. | 2.47 | n.s. | 0.17 | n.s. | <0.01 | n.s. | 0.26 | n.s. | 0.14 | n.s. | 1.19 | n.s. | i. int. | 0.45 | 0.07 | n.s. |
| $Treatment_{Wet}$ | 10.21 | 0.12 ** | 68.38 | 24.05 *** | i. int. | -2.41 | 15.76 | 0.07 *** | 8.95 | 4.43 ** | 57.02 | -8.65 *** | 14.83 | 0.03 *** | i. int. | 3.55 | 0.96 | n.s. |
| CWD | 3.22 | n.s. | 4.63 | 3.31 * | i. int. | -0.7 | 0.86 | n.s. | 0.09 | n.s. | 1.50 | n.s. | 4.52 | 0.01 * | i. int. | -5.13 | 2.13 | n.s. |
| Random effect variances | | | | | | | | | | | | | | | | | | |
| Region:Population | 0.01 (59) | | 28.59 (59) | | 0.14 (59) | | <0.01 (59) | | 6.25 (59) | | 2.16 (59) | | <0.01 (59) | | 15.39 (59) | | 0.59 (59) | |
| Region | <0.01 (17) | | <0.01 (17) | | <0.01 (17) | | <0.01 (17) | | 7.57 (17) | | 3.14 (17) | | <0.01 (17) | | 50.67 (17) | | 5.37 (17) | |
| Residuals | 0.14 (455) | | 0.10 (433) | | 14.68 (450) | | 0.02 (329) | | 184.26 (344) | | 107.89 (360) | | < 0.01 (488) | | 131.73 (442) | | 23.14 (503) | |

Notes: The first column of the table shows the model structure of the maximal models. $\chi$ 2 -values, parameter estimates, and significance levels of fixed effect terms that remained in the minimal adequate models are given. Range and treatment are denoted with the second factor level. Random effect variances of the region and population nested within region are presented with their number of groups in parentheses (i.e., 59 populations, 17 geographical regions). Random effect variances of residuals are presented with the number of observations. n.s., not significant; i. int., not estimated because the main factor was in a significant interaction; significance levels: * p < 0.05; ** p < 0.01; *** p < 0.001.

**Supplementary Table 6.** Details of the linear mixed-effects models investigating how log response ratios of binomial transition traits (germination, survival, flowering), aboveground- and belowground biomass calculated to the experimental drought were related to range (native v. non-native), treatment (wet vs. dry), climatic water deficit (CWD) and their interactions, across 30 native and 29 non-native *C. canadensis* populations.

| | **Performance measures** | | | | | |
|---|---|---|---|---|---|---|
| Fixed effects | Transition variable | | Aboveground biomass | | Belowground biomass | |
| | $\chi^2_{(1)}$ | Estimate | $\chi^2_{(1)}$ | Estimate | $\chi^2_{(1)}$ | Estimate |
| Life $stage_{adult}$ | 65.07 | 0.11*** | 107.38 | 0.44*** | 12.17 | 0.14** |
| $Range_{Non-native}$ : Life $stage_{adult}$ | 0.51 | n.s. | 0.21 | n.s. | 1.95 | n.s. |
| CWD : Life $stage_{adult}$ | 1.53 | n.s. | 0.72 | n.s. | 0.39 | n.s. |
| $Range_{Non-native}$ : CWD : Life $stage_{adult}$ | 3.55 | n.s. | 5.65 | n.s. | 6.05 | 0.16 |

Notes: The first column of the table shows the model structure of the maximal models. χ 2 -values, parameter estimates, and significance levels of fixed effect terms that maximum models are given. Range and treatment are denoted with the second factor level. n.s., not significant; significance levels: * p < 0.05; ** p < 0.01; *** p < 0.001.

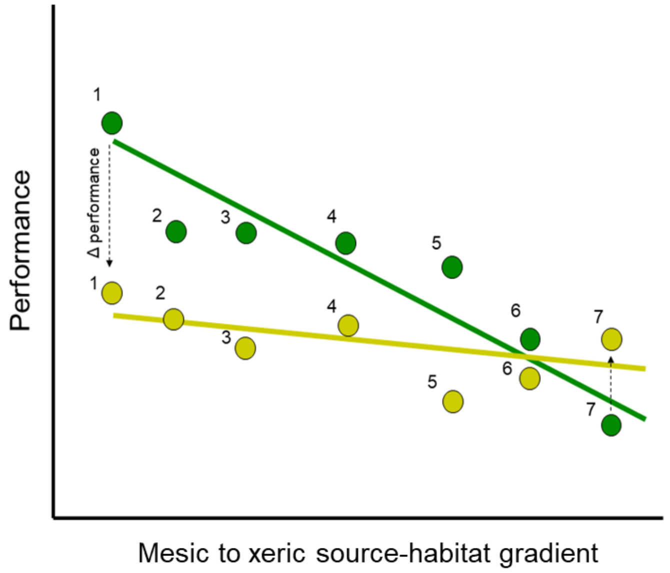


**Supplementary Figure 1.** Concept of clinal variation in the experiment in terms of the drought responses of the performance of populations collected from across a xeric to mesic gradient. Dots represent the populations (numbered) which were sampled on a wide scale from mesic to xeric environment. The figure illustrates a pattern in which plants from the most mesic habitat (population 1) perform much better in wet experimental treatments (green) than in dry experimental treatments (yellow). In contrast, plants from the most xeric habitat (population 7) perform better in dry treatments than in wet treatments. Note that this pattern corresponds to Fig. 1c in the main text (which is simplified for clarity), both showing clinal variation that can be explained by the drought gradient in a way that meets evolutionary expectations. There are many possible permutations of experimental responses along a xeric to mesic gradient (some are illustrated in Figure 1 b-e).

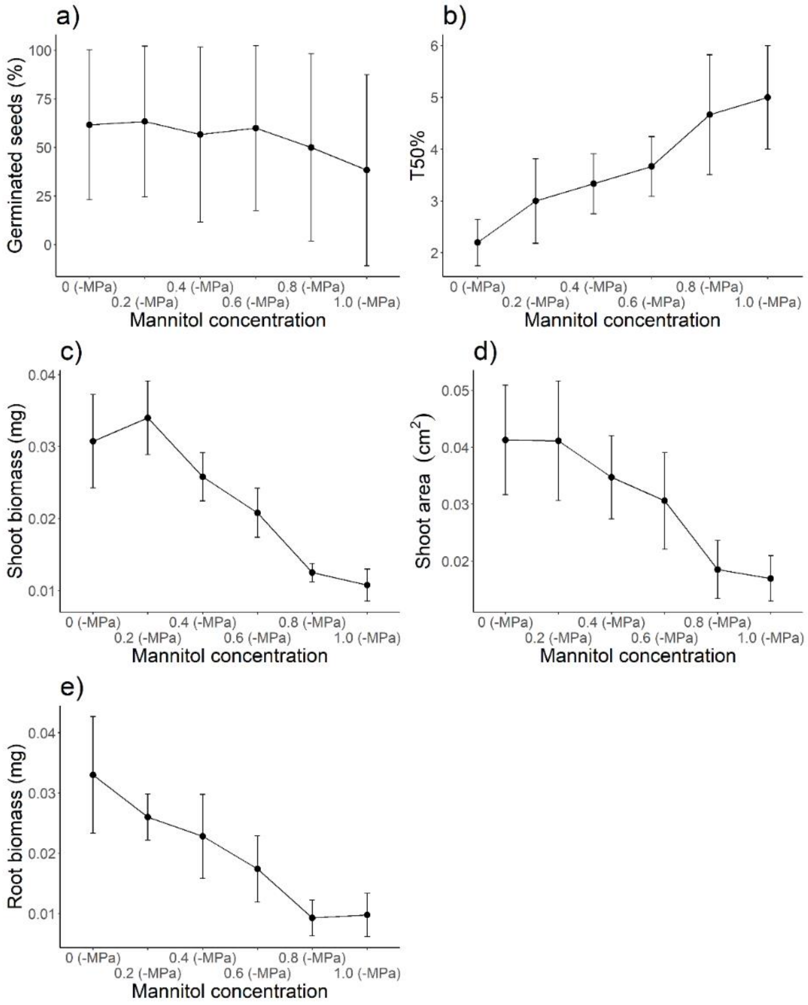


**Supplementary Figure 2.** Results of germination and early seedling traits in a pre-experiment testing six different mannitol concentrations ranging from 0 MPa (tap water) to -1 MPa on six, random populations to determine the effective concentration to be used in the main experiment. Ten seeds from each seed family in 36 Petri dishes (six mannitol concentrations × six seed families) on a filter paper. Applying the same design as in the main experiment the following data were recorded: germinated seeds after 10 days (a), and number of days needed for the 50% germination. Seedling data included (b), dry shoot biomass (c), shoot area (d) and dry root biomass (e) which were assessed for the first germinated seedling of each population 10 days after germination.

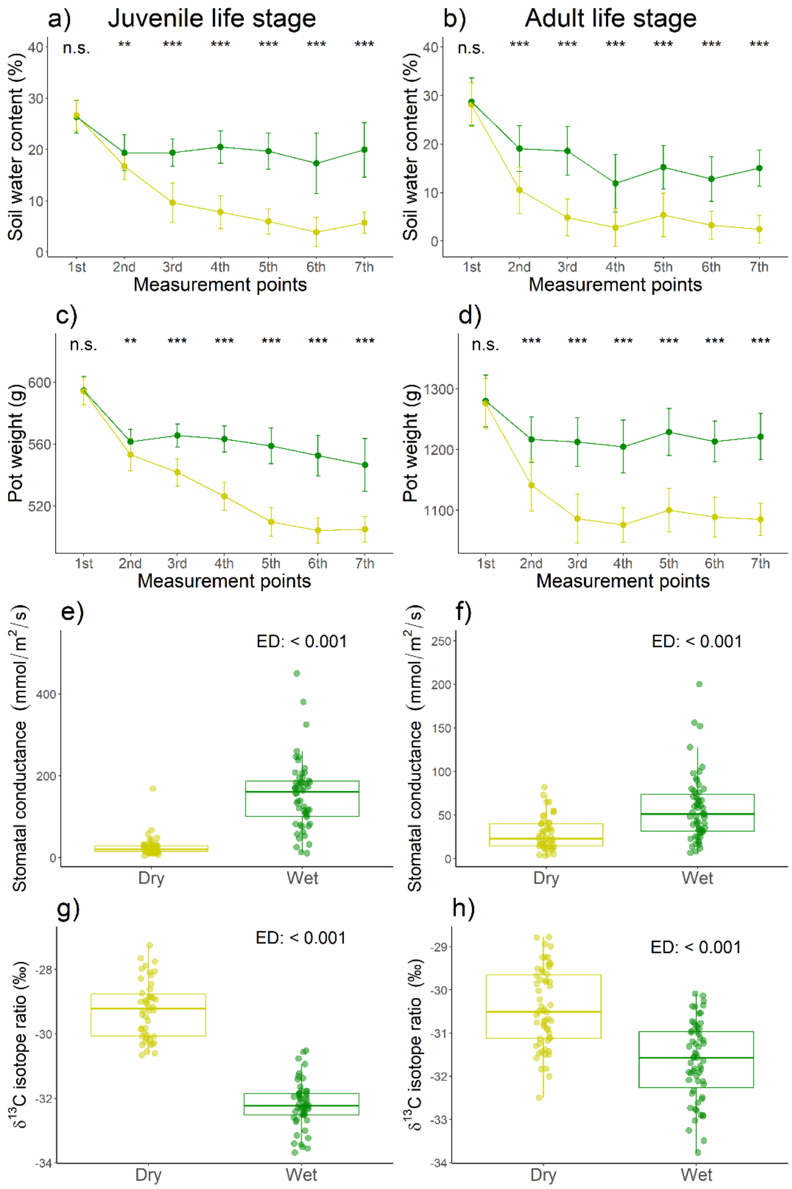


**Supplementary Figure 3.** Measurements of the effects of experimental drought, during the juvenile (a, c, e, g) and adult (b, d, f, h) life stages. Soil moisture content (a - b) and pot weight (c-d) was measured in a weekly basis for juvenile and once in every two weeks for adult plants, on Mondays, before the application of the experimental drought. Stomatal conductance measurements (e – f) were performed before the final harvest. Leaf δ13C isotope ratio (g – h) was determined after the final harvest. Wet treatment (green) refers to the pots that received 100% of their average water loss, while dry treatment (yellow) pots received only 25% every second day. Effect of experimental drought on soil moisture content, pot weight, stomatal conductance and $\delta^{13}C$ was tested using two-sample t-tests. T-values for stomatal conductance and $\delta^{13}C$ were collected: stomatal conductance at juvenile life stage ($t = 10.55$), and adult life stage ($t = 4.81$), $\delta^{13}C$ at juvenile life stage ($t = -13.57$), and adult life stage ($t = -7.31$). For soil water content and pot weight, asterisks indicate significant differences between wet vs. dry treated plants at the different measurement points through time: *, $P < 0.05$; **, $P < 0.01$; ***, $P < = 0.001$; n.s., non-significant.

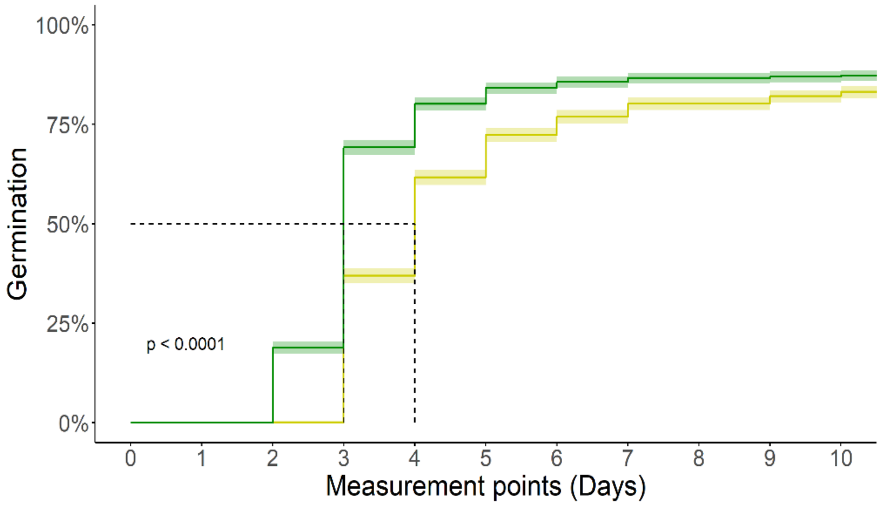


**Supplementary Figure 4.** Kaplan-Meier plot for germination success *Conyza canadensis* across the 10 days long experiment. Effect of experimental drought (wet (green) vs. dry (yellow)) on germination was tested. The dashed line refers to the timepoint when 50% of all seeds germinated (including those that finally never germinated). Confidence intervals for each measurement point are visualized. Details on the models and results can be found in Suppl. Table S4.

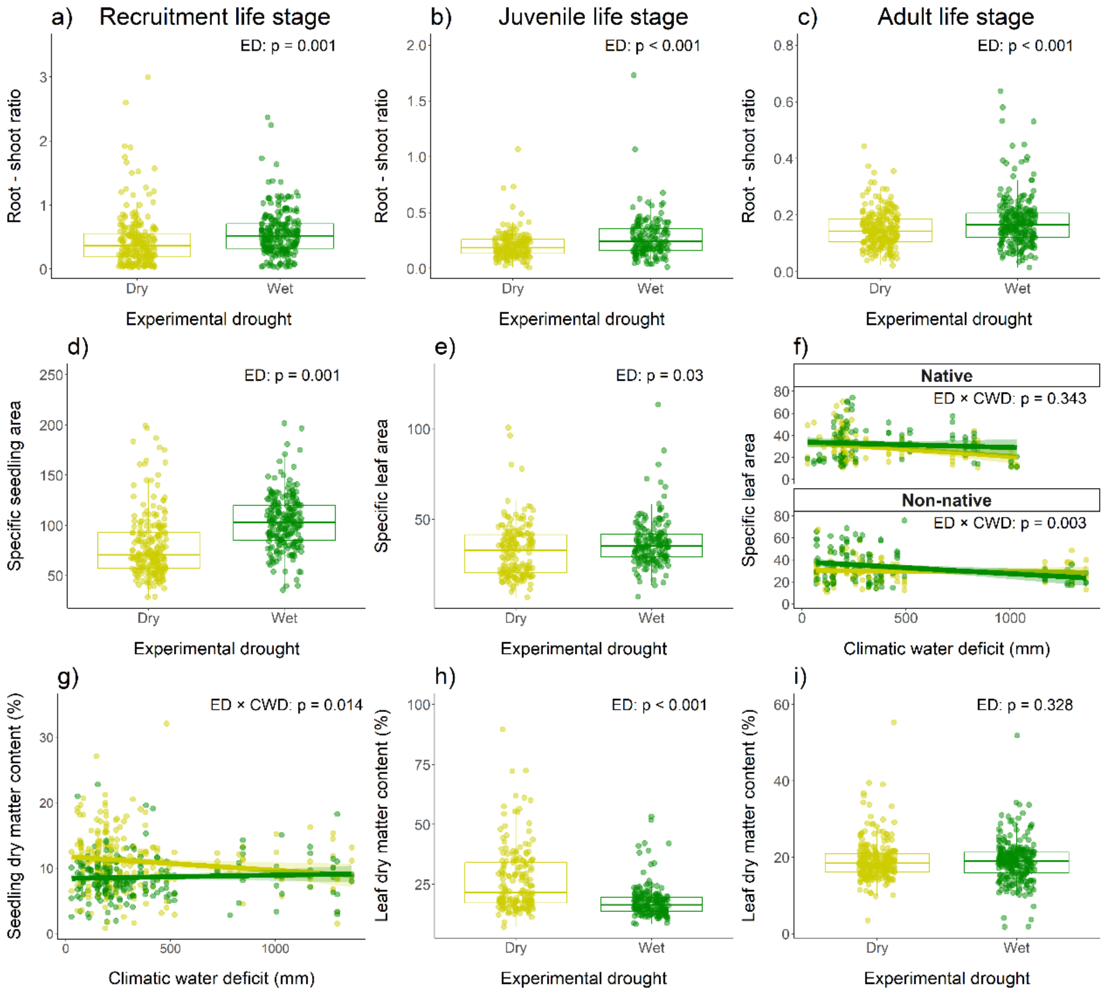


**Supplementary Figure 5.** Interactive effects of experimental drought (ED), climatic water deficit (CWD) and range-affiliation (native vs. non-native) on plant functional traits for root-shoot ratio (a-c), specific leaf area (d-f) and leaf dry matter content (g-i). Functional traits were recorded across three experiments focusing on the following life stages: germination and seedling stage (a, d, g), juvenile stage (b, e, h) and adult stage (c, f, i). The figures represent the results of the minimum adequate models (i.e., including only significant terms), except for the response variables where none of the three tested explanatory factors or their interactions had a significant effect (here: leaf dry matter content at adult stage (i)). Details on the models and results can be found in Supplementary Table 5.

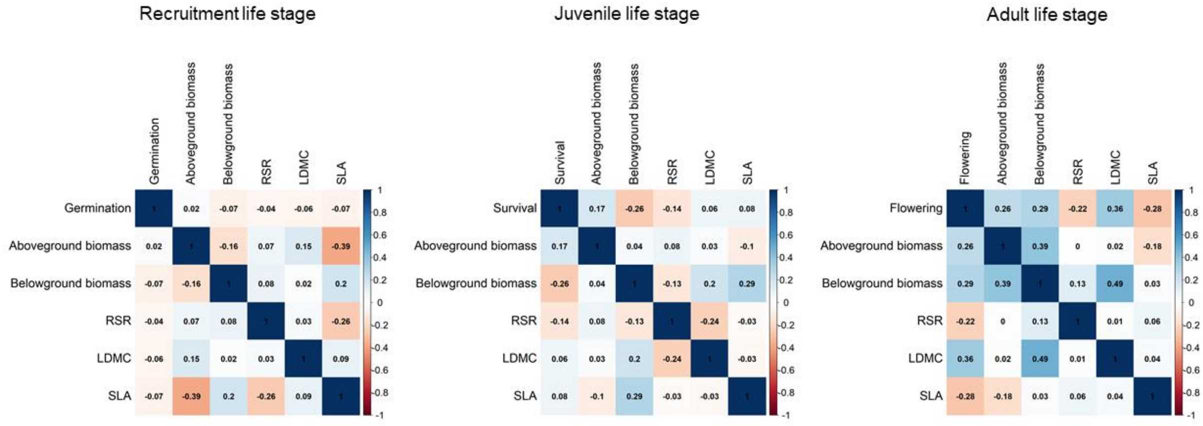


**Supplementary Figure 6.** Details of the analyses investigating the correlation among the calculated log response ratios of performance (germination, survival, flowering, aboveground- and belowground biomass) and functional traits (root-shoot ratio, specific leaf area and leaf dry matter content) in response to the applied experimental drought, across life stages, in 30 native and 29 non-native *C. canadensis* populations. Pearson correlation coefficients are presented for each pair-wise correlation (see also legend).

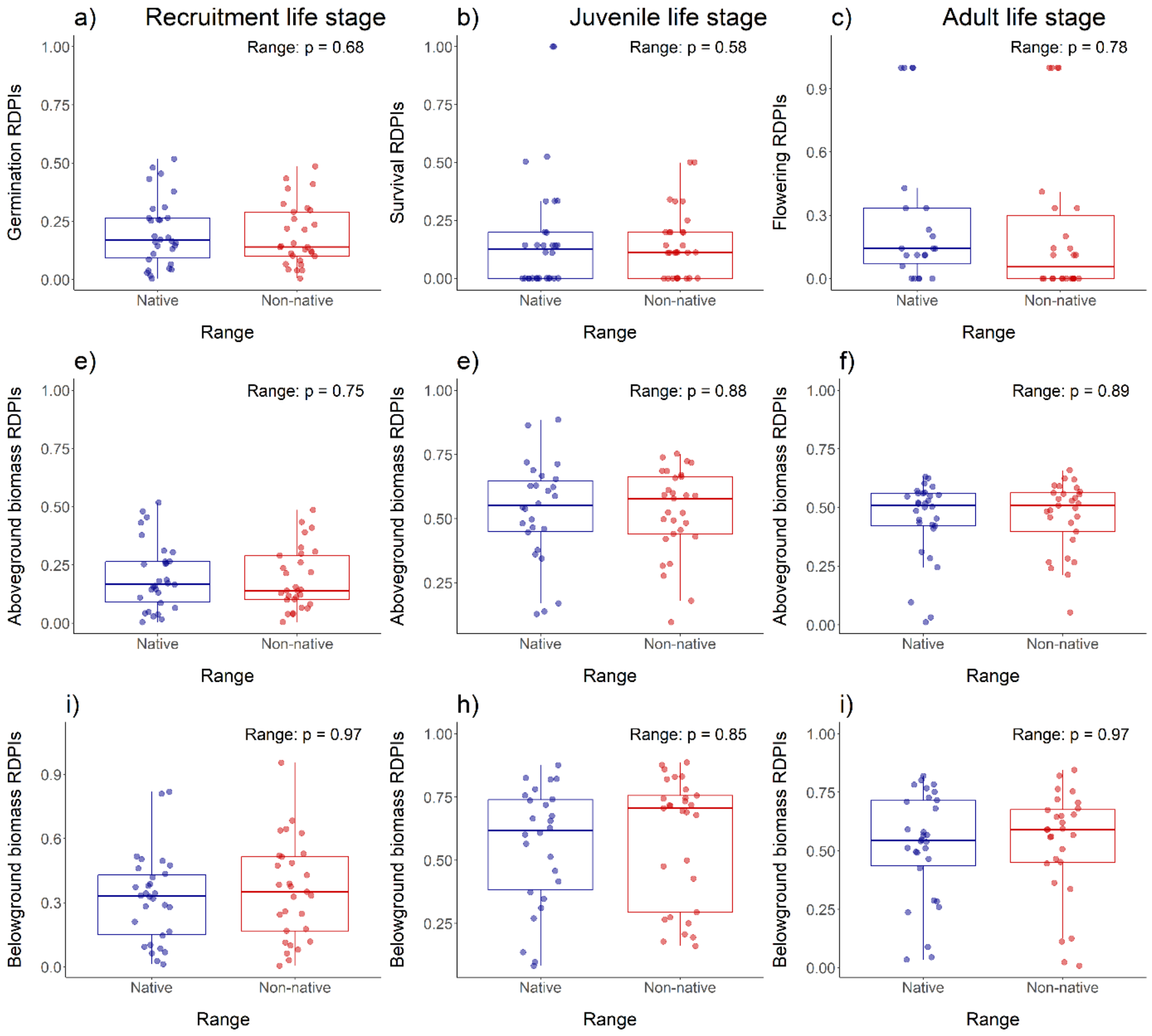


**Supplementary Figure 7.** Effects of range-affiliation (native vs. non-native) on plasticity of plant performance for germination (a), survival (b) and flowering (c), aboveground biomasses (d-f) and belowground biomasses (g-i). Plasticity refers to as the plastic responses to experimental drought (i.e., relative plasticity indexes; RDPIs). Performance measures were recorded across three experiments focusing on the following life stages: recruitment life stage (a, d, g), juvenile life stage (b, e, h) and adult life stage (c, f, i).

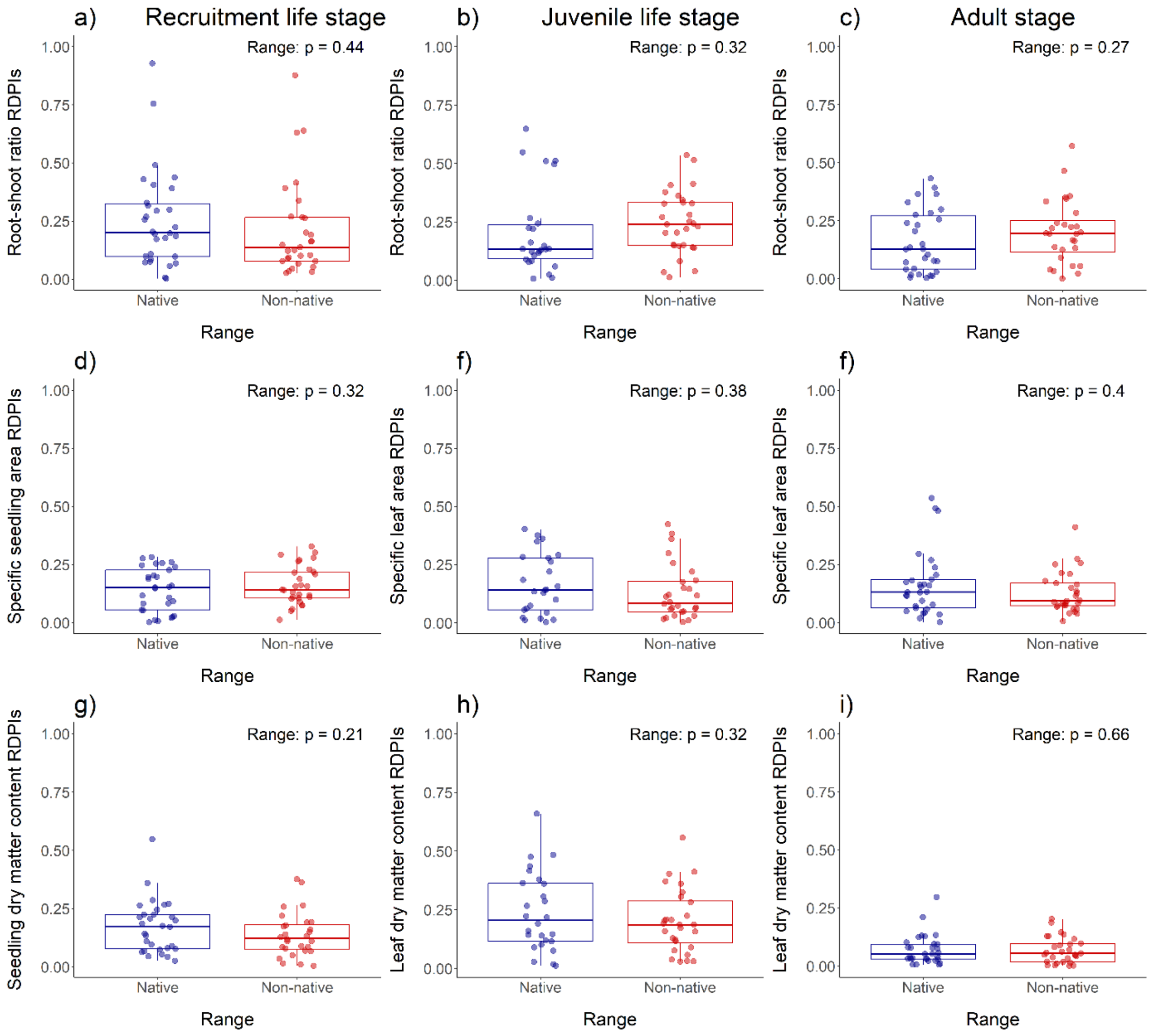


**Supplementary Figure 8.** Effects of range-affiliation (native vs. non-native) on plasticity of plant functional traits for root-shoot ratio (a-c), specific leaf area (d-f) and leaf dry matter content (g-i). Plasticity refers to as the plastic responses to experimental drought (i.e., relative plasticity indexes; RDPIs). Functional traits were recorded across three experiments focusing on the following life stages: recruitment life stage (a, d, g), juvenile life stage (b, e, h) and adult life stage (c, f, i).
